# Cryo-EM structure analysis of phage ΦXacm4-11 that infects the phytopathogen *Xanthomonas citri*

**DOI:** 10.64898/2026.02.16.706137

**Authors:** Chuck Shaker Farah, Guilherme Oliveira Silva, Edgar Enrique Llontop, Alexandre Cassago, German Dunger, Jeffrey B. Jones, João Carlos Setubal, Aline Maria da Silva, Rodrigo Portugal, Germán Gustavo Sgro

**Affiliations:** Departamento de Bioquímica, Instituto de Química, Universidade de São Paulo, São Paulo, SP, Brazil; Departamento de Ciências Biomoleculares, Faculdade de Ciências Farmacêuticas de Ribeirão Preto, Universidade de São Paulo, Ribeirão Preto, SP, Brazil; Laboratório Nacional de Nanotecnologia, Centro Nacional de Pesquisa em Energia e Materiais, Campinas, SP, Brazil; Departamento de Producción Animal, Facultad de Ciencias Agrarias, Universidad Nacional del Litoral, Esperanza, SF, Argentina; Department of Plant Pathology, University of Florida, Gainesville, Florida, USA

**Keywords:** *podoviridae*, Single particle analysis cryo-electron microscopy, bacteriophage structure, *Xanthomonas citri*

## Abstract

Very few bacteriophages that infect *Xanthomonas* species have been characterized genetically and only one 3D structure, the capsid of a siphovirus that infects the phytopathogen *Xanthomonas citri*, has been determined at high resolution. This study presents the annotated DNA sequence and detailed structural analysis of ΦXacm4-11, a podovirus that infects *Xanthomonas citri*, shedding light on its unique architecture and functional attributes, providing insights into the molecular mechanisms underlying host recognition and infection. Annotation of the genome revealed conserved features among related phages, but also distinct genetic elements that may contribute to ΦXacm4-11’s specificity toward *X. citri.* Genes associated with host recognition and infection were identified, including the genes potentially coding for the receptor-binding proteins (RBPs) at the tail fibre tip, offering insights into their role in bacterial attachment. Using high-resolution cryo-electron microscopy, we resolved the architecture of the mature, pre-released virion, revealing a T7-like head-tail assembly with a well-defined portal-tail complex embedded at a unique fivefold vertex. Our findings provide a detailed view of the structural and functional components of ΦXacm4-11, furthering our understanding of its molecular interactions with *X. citri* and its potential application in phage therapy against phytopathogens.

**SIGNIFICANCE STATEMENT:** Bacteriophages are increasingly recognized as powerful tools to control bacterial pathogens in medicine and agriculture, yet the structural basis of host recognition and genome delivery remains poorly understood for most phages. Here, we present a comprehensive structural and functional analysis of ΦXacm4-11, a podovirus that infects the plant pathogen *Xanthomonas citri*. By combining genome annotation, proteomics, and high-resolution cryo-electron microscopy, we reveal the complete architecture of the mature virion and its specialized portal-tail machinery. Our results show how this short-tailed phage deploys an internal injection device to penetrate the bacterial cell envelope and highlight structural features linked to type IV pilus-dependent infection. These findings provide insights into phage entry mechanisms and establish ΦXacm4-11 as a model for engineering biocontrol strategies.

## INTRODUCTION

Phages are perhaps the most abundant replicating structures on earth [1–3] and, in light of the inevitable emergence of multidrug resistant strains of pathogenic bacteria [4], there is an ever growing interest in engineering phages to efficiently target specific bacterial populations [5,6]. They have been employed to combat bacterial infections in humans [7–9], livestock [10,11] and agricultural crops [12,13], demonstrating plasticity and broad potential for biotechnological approaches. Phages use a variety of mechanisms to attach to and enter host bacterial cells [14]. In Gram-negative bacteria they mostly target O-antigens (smooth LPS), core oligosaccharides (rough LPS), surface-exposed proteins (e.g. LamB, TolC, FepA, OmpA, OmpC and OmpF), flagella, pili and capsule; while in Gram-positive bacteria they mostly target teichoic acids, surface-exposed proteins (e.g. GamR and YueB) and peptidoglycan structures (PG) [14]. Type IV pilus (T4P)-dependent phage infection is very common in many bacterial species, including Xanthomonadaceae species. For example, *X. citri* infection by the ΦXacm4-11 podovirus [15,16] and Cf filamentous phage [17,18], *Xylella fastidiosa* strain Temecula 1 and *Xanthomonas* sp. strain EC-12 infections by the siphophages Sano and Salvo and the podophages Prado and Paz [19], and others [20] are all dependent on bacterial T4P.

The literature on bacteriophage biology has, to date, primarily focused on well-characterised representatives (e.g. enterobacteria T4, λ and T7 phages). However, the vast genomic diversity among bacteriophages suggests a reservoir of untapped biological insights waiting to be uncovered. *Xanthomonas* species infect several hundred different plant hosts, including many economically important crops. Bacteriophages that infect *Xanthomonas* species are therefore of interest as potential biocontrol agents. Many bacteriophages that infect *Xanthomonas* species in general [19,21–24] and *X. citri* strains in particular [16,25–27], have been isolated and characterised for their microbiological features. However, although a handful of their genomes have been sequenced [28–35] none have been studied using multidisciplinary structural/functional approaches and no high resolution structural information is available for bacteriophages that infect Xanthomonas bacteria.

The bacteriophage ΦXacm4-11 was first isolated from citrus tissue as infecting *X. alfalfae* subsp. *citrumelonis* strain 36 (Xacm36) in Florida, and has been shown to be able to infect a large number of *X. citrumelo* and *X. citri* isolates from around the world [16]. Phage ΦXacm4-11 forms clear plaques on a lawn of the reference *X. citri* strain 306 (Xac306) [36], for which our group previously demonstrated that the infection is dependent on a functional Type IV pilus (T4P) [15,37]. Bacterial T4P are long, flexible surface filaments that consist of helical polymers of both major and minor pilin subunits. The ability of these pili to polymerize, attach and depolymerize mediates several bacterial behaviours, including twitching motility, surface adhesion, pathogenicity, natural transformation, escape from immune system defence mechanisms, and biofilm formation [15].

In this study, we present the complete annotated genome sequence of bacteriophage ΦXacm4-11. The genome reveals key genes involved in phage assembly, DNA packaging, and host recognition, providing insights into the molecular mechanisms underlying its specificity for *X. citri*. We also describe the high-resolution structure of the mature, pre-released virion obtained through cryo-electron microscopy, highlighting its unique architectural features, including the capsid, tail complex, and potential receptor-binding proteins. These structural details further enhance our understanding of phage-host interactions and the potential application of ΦXacm4-11 in phage-based bacterial control.

## RESULTS AND DISCUSSION

### Microbiological characterization of ΦXacm4-11

To determine key parameters of the phage replication cycle, we conducted adsorption and one-step growth assays using ΦXacm4-11 and *X. citri* strain 306 as host (**Figure 1A** and **Figure S1**). The assays revealed a latent period of 60 minutes and a burst size of 299 particles released per infected cell, indicating an efficient lytic replication cycle. Negatively stained electron microscopy images of purified phage particles showed a regular icosahedral capsid ∼70 nm in diameter connected to a ∼22 nm short tail (**Figure 1B**), suggesting that ΦXacm4-11 belongs to the *Podoviridae* family of bacteriophages. When examined together, multiple ΦXacm4-11 particles displayed a preferential orientation toward previously purified *X. citri* strain 306 T4P in which the tails were employed as points of attachment (**Figure 1B**). This confirms that the ΦXacm4-11 particle has receptors specific for T4P epitopes, and is consistent with previous demonstrations that *X. citri* strain 306 T4P is required for ΦXacm4-11 infection [15,37].

**Figure 1.**
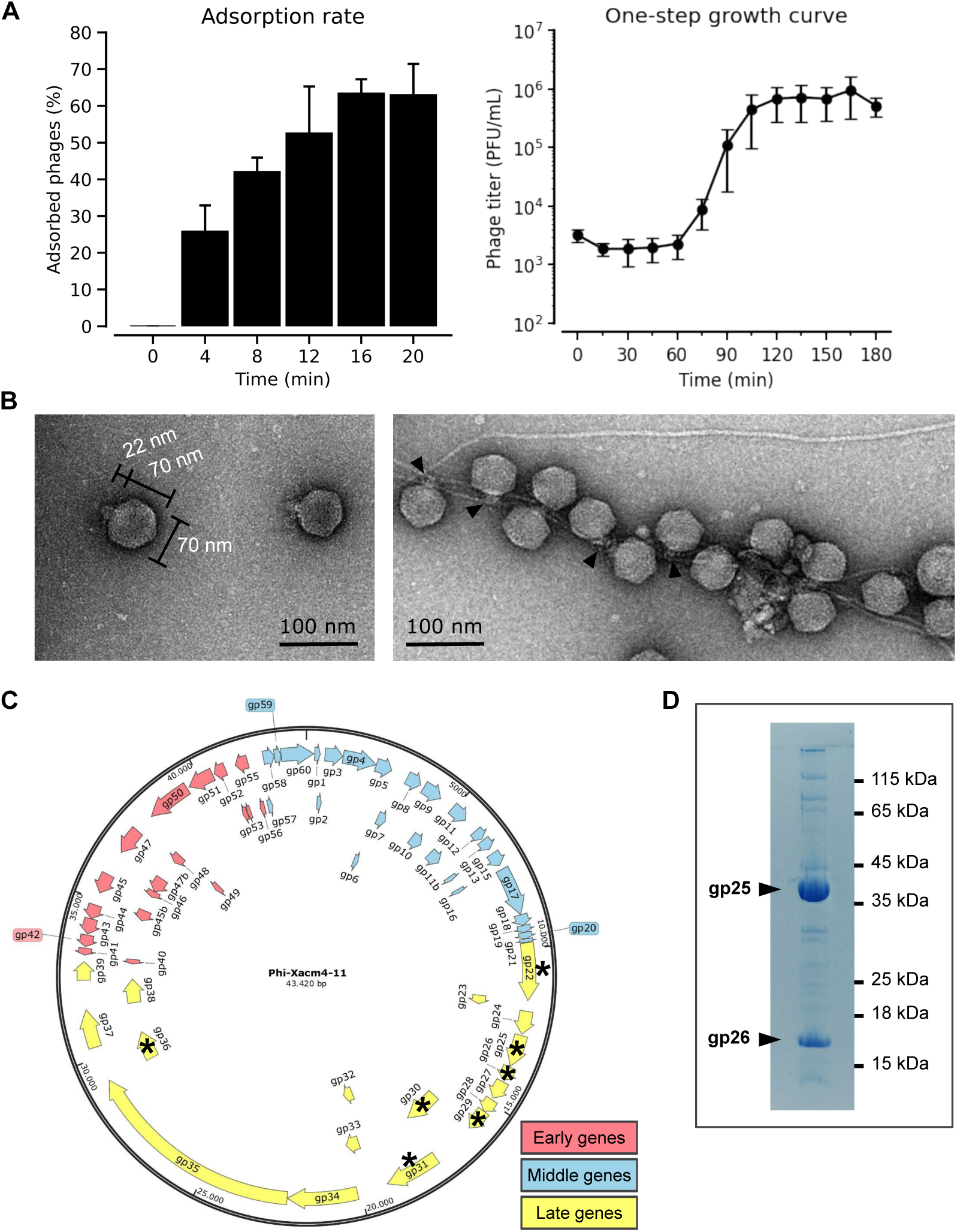
Microbiological and genomic characterization of ΦXacm4-11. **(A)** *Left:* Adsorption kinetics of ΦXacm4-11 to *Xanthomonas citri*. Phage particles were mixed with host cells and incubated at room temperature. At the indicated time points, samples were taken, and the title of non-adsorbed (free) phages was determined by plaque assay after removal of bacterial cells. The fraction of adsorbed phages was calculated relative to the initial phage input. *Right:* One-step growth curve of ΦXacm4-11 used to determine key parameters of the phage replication cycle. The latent period (∼60 min) and burst phase (∼60-105 min) were estimated from the growth curve, and the burst size was calculated as 299 ± 110. Data were obtained from four independent biological replicates. Viral titers were log_10_-transformed prior to analysis. Mean values and SEM were calculated in logarithmic space and are shown as error bars. **(B)** Negative-stain transmission electron micrographs of ΦXacm4-11 particles. *Left*: isolated virions exhibiting an icosahedral capsid and a short tail, consistent with podovirus morphology. *Right:* virions interacting with purified *X. citri* type IV pilus filaments (black arrows). **(C)** Schematic representation of the ΦXacm4-11 genome showing predicted open reading frames (ORFs) following comparative genomic analysis. ORFs are displayed as arrows according to the coding strand and colored by functional category: involved in taking over the host cell, shutting down host DNA/RNA/protein synthesis, and modifying host RNA polymerase (early genes, red); responsible for DNA replication, repair, and recombination (middle genes, blue); and encoding structural components for the phage head and tail, as well as proteins for cell lysis (late genes, yellow). Bold asterisks indicate ORFs for which protein structures were modelled in the cryo-EM maps. For more information, refer to **Table S1**. **(D)** 16% Tricine SDS-PAGE analysis of CsCl-purified ΦXacm4-11 virion particles. Prominent bands correspond to the major capsid protein (MCP, gp25, 36.4 kDa) and the cement protein (CP, gp26, 15.4 kDa).

### Sequencing and annotation of the ΦXacm4-11 genome

The ΦXacm4-11 genome was sequenced by pair-ended Illumina technology and assembled into a single contig 43,420 bp long (**Figure 1C**). The randomly fragmented NGS data were analyzed using the PhageTerm software [38] to predict the bacteriophage physical termini and DNA packaging mode (**Figure S2A**). This analysis identified 1,108 bp direct terminal repeats (DTRs), consistent with a T5-like DNA packaging mechanism (long DTR). The presence of fixed termini at positions 9,535 and 10,642, together with the non-permuted genome organization, indicates a linear DNA molecule with direct terminal redundancy. Although limited in spatial resolution, complementary PCR assays using primers targeting regions adjacent to the contig predicted physical termini generated amplification products consistent with a linear genome organization inside the virion (**Figure S2B**).

Bioinformatic analysis of the nucleotide and translated sequences (**File S1** and **File S2**) showed that the ΦXacm4-11 genome is most similar to that of *X. citri* phage Cp2 [28] (92% nucleotide identity over 86% coverage of Cp2), placing it within the *Podoviridae* family, and confirming our morphologic classification using electron microscopy.

The ΦXacm4-11 DNA chromosome codes for 63 open reading frames (ORFs) and corresponding gene products (gps) (**Figure 1C** and **Table S1**), which were numbered in a manner that partially follows the annotation of the Cp2 genome, in spite of the fact that the latter has only 40 annotated ORFs [28]. The genes can be divided into three main clusters based on collinearity and similarities with other podoviridae genomes, with the class I genes (gp40 to gp56) being coded on the opposite strand of that of the class II (gp1 to gp21 and gp57 to gp60) and class III genes (gp22 to gp39).

The class III gene cluster codes for structural proteins making up the capsid (gp25 and gp26), the tail (gp22, gp29, gp30 and gp31) and tail fibres (gp36 to gp39) as well as other proteins of unknown function. The class I and II gene clusters mostly code for hypothetical proteins and proteins involved in DNA processing. The class I cluster, expected to code for early function proteins, includes an exonuclease (gp45), two endonucleases (gp45b and gp47b), and two single-stranded DNA (ssDNA) binding proteins (gp44 and gp47), with gp44 and gp45 showing similarity to the anti-CRISPR factors Vcrx091 and Vcrx93, respectively, from the IncC conjugative plasmid pVCR94 [39,40]. The class II proteins include two endonucleases (gp5 and gp11b), a recA-dependent nuclease (gp7), a crossover junction resolvase (gp8), a lysozyme-like protein (gp9) and two terminase-related proteins (gp15 and gp17). No RNA polymerase or tRNA genes were identified.

The protein content of purified phage particles was analysed by SDS-PAGE separation (**Figure 1D**), providing an overview of the relative abundance of protein bands. Shotgun mass spectroscopy proteomics of purified phage identified 47 of the 63 predicted gene products (**Table 1**). Proteomics was also performed on viral proteins separated by SDS-PAGE (in-gel proteomics). Combined, thirty-four of these gene products were identified by at least two peptide hits and greater than 10% coverage. In this reduced list, 15 of the 18 class III (83.3%) gene products were identified (14 had over 40% coverage). On the other hand, using the same criteria of at least two peptide hits and greater than 10% coverage, class I (8 out of 19; 42.1%) and class II proteins (11 out of 26; 42.3%) were much less abundant in the mature particles.

**Table 1.** List of identified gene products (gps) in purified ΦXacm4-11 by shotgun proteomics.

| Gene product | Predicted function | Number of AAs | Coverage (%) | Number of peptides | MW (kDa) |
| --- | --- | --- | --- | --- | --- |
| gp1 | Hypothetical protein | 55 | 21.82 | 1 | 6.3 |
| gp2 | Hypothetical protein | 56 | 39.29 | 2 | 6.3 |
| gp3 | Hypothetical protein | 184 | 42.39 | 3 | 20.9 |
| gp4 | Hypothetical protein | 349 | 20.06 | 4 | 38.9 |
| gp5 | HNH Endonuclease | 168 |  |  | 19.2 |
| gp6 | Hypothetical protein | 82 |  |  | 9.3 |
| gp7 | RecA-dependent nuclease | 111 |  |  | 12.5 |
| gp8 | RusA endodeoxynuclease | 156 | 7.69 | 1 | 16.9 |
| gp9 | Lysozyme-like superfamily | 206 | 59.22 | 10 | 22.1 |
| * gp10 | Hypothetical protein | 153 | 13.73 | 1 | 15.9 |
| * gp11 | Hypothetical protein | 185 | 7.57 | 1 | 20.3 |
| gp11b | HNH endonuclease | 169 |  |  | 18.6 |
| gp12 | Hypothetical protein | 104 |  |  | 11.5 |
| gp13 | Hypothetical protein | 114 |  |  | 12.0 |
| gp14 | Hypothetical protein | 54 |  |  | 5.5 |
| gp15 | Terminase small subunit | 143 | 18.88 | 2 | 16.0 |
| gp16 | Hypothetical protein | 47 |  |  | 5.5 |
| * gp17 | Terminase large subunit | 508 | 23.03 | 7 | 56.2 |
| * gp18 | Hypothetical protein | 112 | 94.64 | 11 | 12.0 |
| gp19 | Hypothetical protein | 71 | 42.25 | 2 | 7.6 |
| * gp20 | Hypothetical protein | 66 | 75.76 | 5 | 6.8 |
| gp21 | Hypothetical protein | 37 |  |  | 4.2 |
| * gp22 | **Portal protein** | 598 | 88.29 | 50 | 65.6 |
| gp23 | Hypothetical protein | 113 | 45.13 | 2 | 13.0 |
| * gp24 | Putative protease | 241 | 42.32 | 8 | 25.0 |
| * gp25 | **Major capsid protein** | 337 | 77.45 | 30 | 36.4 |
| * gp26 | **Cement protein** | 148 | 75.00 | 11 | 15.4 |
| gp27 | Hypothetical protein | 196 | 5.10 | 1 | 19.9 |
| gp28 | Endonuclease | 143 | 4.90 | 1 | 15.3 |
| * gp29 | **Adaptor protein** | 227 | 77.53 | 15 | 24.8 |
| * gp30 | **Tail tubular protein** | 463 | 75.81 | 28 | 49.6 |
| * gp31 | **Nozzle protein** | 570 | 71.75 | 34 | 61.9 |
| gp32 | Internal virion protein | 158 | 29.11 | 1 | 17.8 |
| * gp33 | Internal virion protein | 167 | 76.65 | 13 | 17.0 |
| * gp34 | Lysozyme-like protein / Endolysin | 730 | 83.29 | 74 | 79.1 |
| * gp35 | Hypothetical protein | 2300 | 78.91 | 183 | 248.9 |
| * gp36 | **Tail fiber protein** | 369 | 88.08 | 23 | 40.1 |
| * gp37 | Tail fiber protein | 399 | 31.83 | 7 | 42.2 |
| * gp38 | Tail fiber protein | 310 | 44.19 | 5 | 32.9 |
| * gp39 | Tail fiber protein | 191 | 53.93 | 12 | 21.6 |
| gp40 | Hypothetical protein | 59 | 77.97 | 4 | 6.4 |
| gp41 | Hypothetical protein | 71 | 84.51 | 5 | 8.0 |
| gp42 | Hypothetical protein | 120 |  |  | 12.9 |
| * gp43 | Hypothetical protein | 161 | 8.07 | 2 | 17.7 |
| gp44 | Anti-CRISPR Vcrx091 protein | 145 |  |  | 16.4 |
| gp45b | HNH endonuclease | 136 |  |  | 15.6 |
| * gp45 | Exonuclease | 207 | 40.10 | 6 | 23.3 |
| * gp46 | Hypothetical protein | 80 | 26.25 | 2 | 8.6 |
| gp47b | Homing endonuclease | 194 |  |  | 21.7 |

**Table 1 (cont.)** | List of identified gene products (gps) in purified ΦXacm4-11 by shotgun proteomics.
| Gene product | Predicted function | Number of AAs | Coverage (%) | Number of peptides | MW (kDa) |
| --- | --- | --- | --- | --- | --- |
| gp47 | ssDNA-binding protein | 279 | 7.53 | 2 | 30.3 |
| gp48 | Hypothetical protein | 116 | 56.03 | 5 | 13.1 |
| gp49 | Hypothetical protein | 91 |  |  | 10.0 |
| * gp50 | Hypothetical protein | 470 | 20.21 | 5 | 52.6 |
| * gp51 | Hypothetical protein | 274 | 45.26 | 10 | 29.4 |
| gp52 | Hypothetical protein | 124 | 54.84 | 3 | 13.1 |
| gp53 | Hypothetical protein | 60 |  |  | 7.1 |
| gp54 | Hypothetical protein | 49 |  |  | 5.3 |
| gp55 | Hypothetical protein | 132 | 5.30 | 1 | 14.3 |
| gp56 | Hypothetical protein | 60 | 10.00 | 1 | 6.9 |
| gp57 | Hypothetical protein | 63 | 61.90 | 5 | 7.1 |
| * gp58 | Hypothetical protein | 115 | 10.43 | 1 | 12.5 |
| gp59 | Hypothetical protein | 68 | 50.00 | 1 | 7.5 |
| gp60 | Hypothetical protein | 339 | 48.95 | 4 | 37.3 |

\* Proteins also found by SDS-PAGE in-gel proteomics.

### Cryo-EM structure of ΦXacm4-11 capsid

Using datasets acquired on a FEI Titan Krios 300 kV microscope, we successfully processed and reconstructed the ΦXacm4-11 viral particle *de novo* by single-particle analysis in RELION [41], without applying any symmetry constraints (**Figure S3**). Briefly, after several rounds of 2D and 3D classifications, a subset of 65,055 particles yielded a map with 4.11 Å resolution from which we identified many features in considerable detail of the phage portal and tail, embedded in the 5-fold symmetrical special vertex of the icosahedral capsid (**Figure 2**, **Figure S4A-B**). The asymmetric reconstruction revealed not only detailed features of the tail architecture but also multiple internal structural elements, including the nozzle, adaptor, portal, DNA-protein injection complex, and core protein complex. The slice cut of the C1 map also showed 6-8 concentric layers of double-stranded DNA density within the capsid (**Figure S4A**). These layers are ∼2.4 nm apart (center to center) as expected for closely-packed DNA double helices. The outer layer of DNA strands is closely juxtaposed with the inner capsid lumen formed mainly by the bottom surface of the major capsid protein (**Figure S4C**). The resolution of the structure does not allow us to model the DNA molecule or analyze specific protein-DNA interactions.

**Figure 2.**
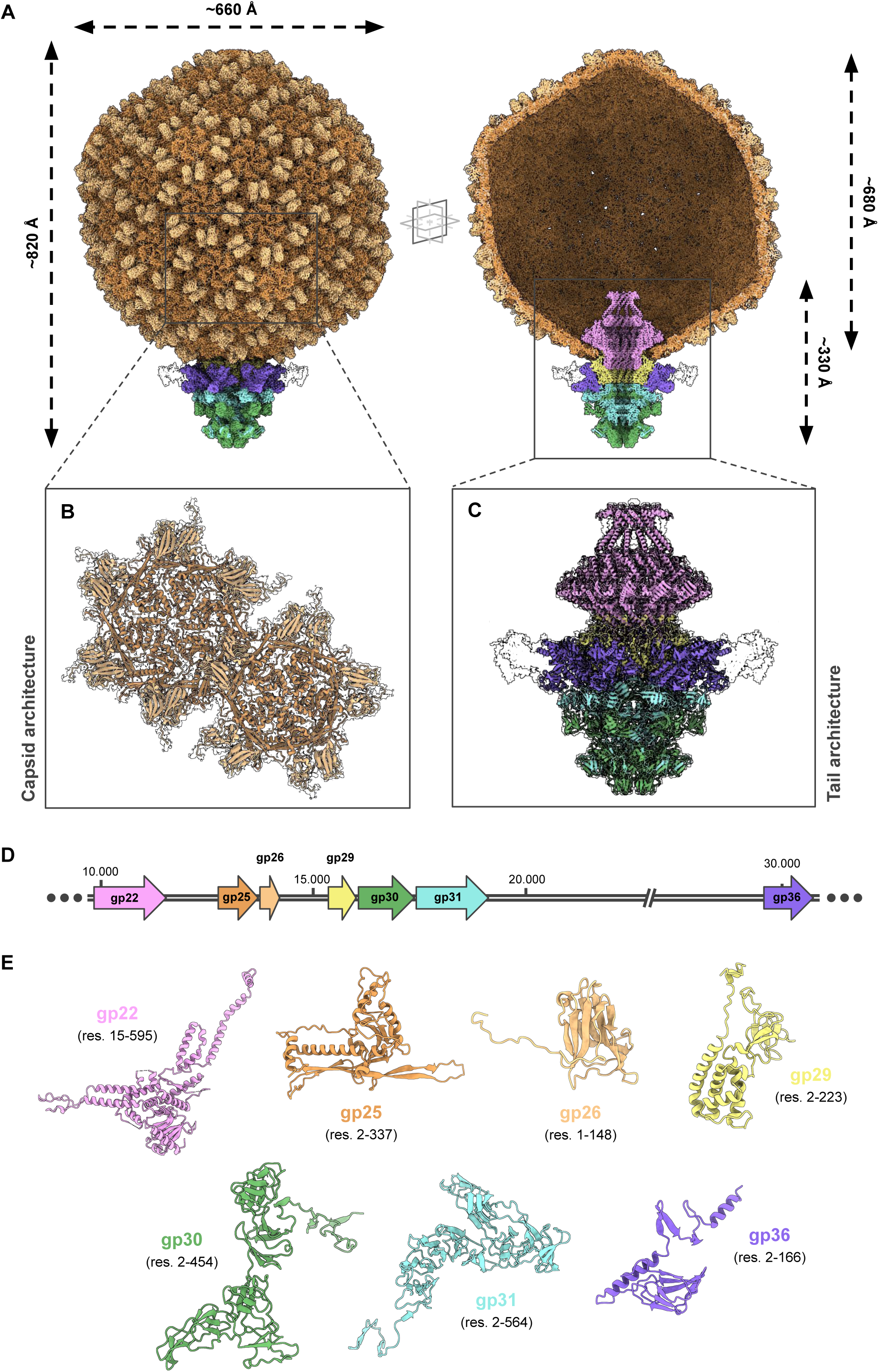
Structural architecture of phage ΦXacm4-11 solved by cryo-EM. **(A)** *Left*: Reconstruction of the full virion, showing its external surface. *Right*: Longitudinal section along the phage axis, revealing internal densities and tail organization. Overall virion dimensions are indicated. **(B)** Magnified view of the capsid highlighting the arrangement of capsomers in distinguishable hexons and pentons, and the spatial distribution of capsid-associated decoration proteins. **(C)** Magnified view of the tail, presented as a cryo-EM map with fitted atomic models, illustrating the architecture of the portal complex and basal structures. **(D)** Cluster of structural gene products of ΦXacm4-11, highlighting the ORFs of proteins that were modelled in this study. **(E)** Atomic models of the resolved structural proteins, shown individually and colour-coded as they appear in the preceding panels, including the major capsid protein (gp25), cement protein (gp26), portal (gp22), adaptor (gp29), two nozzle subunits (gp30 and gp31) and a tail fibre subunit (gp36).

Once the asymmetric map of the capsid was obtained, we focused on achieving higher-resolution reconstructions and structural models for the ΦXacm4-11 virions. Using 69,590 particles extracted from the same datasets, we performed icosahedral (symmetry-imposed) single-particle reconstruction in RELION [41], yielding a 3.16-Å-resolution map of the ΦXacm4-11 capsid (**Figure 3A-B**, **Figure S5A,C** and **Table S2**). The near-atomic density map (**Figure 3B**) shows that the phage particle possesses a *T* = 7 (*laevo*) shell containing 60 asymmetric units (ASU), each of which in turn is made up of seven copies of the major capsid protein (MCP, gp25) and seven copies of the cement protein (CP, gp26). We then built and iteratively refined an entire capsid asymmetric unit (MCP subunits A-G and CP subunits H-N) and a complete icosahedral virion capsid until good model-to-map correlation results were obtained (**Figure 3C-F** and **Table S2**). The asymmetric unit consists of one pseudo-symmetric hexon (subunits A-F) plus one fifth of a symmetric penton (subunit G). The CP subunits are found as upright pairs at the hexon-hexon and hexon-penton interfaces (**Table S3**) and decorate the external surface of the capsid (**Figure 3A,E**) in an organisation similar to that previously observed for the bacteriophages BPP-1 from *Bordetella pertussis* [42] and ε15 from *Salmonella anatum* [43]. The capsid has a thickness of ∼40-60 Å depending on the region, and a maximum diameter of ∼680 Å, as measured between two opposite 5-fold vertices.

**Figure 3.**
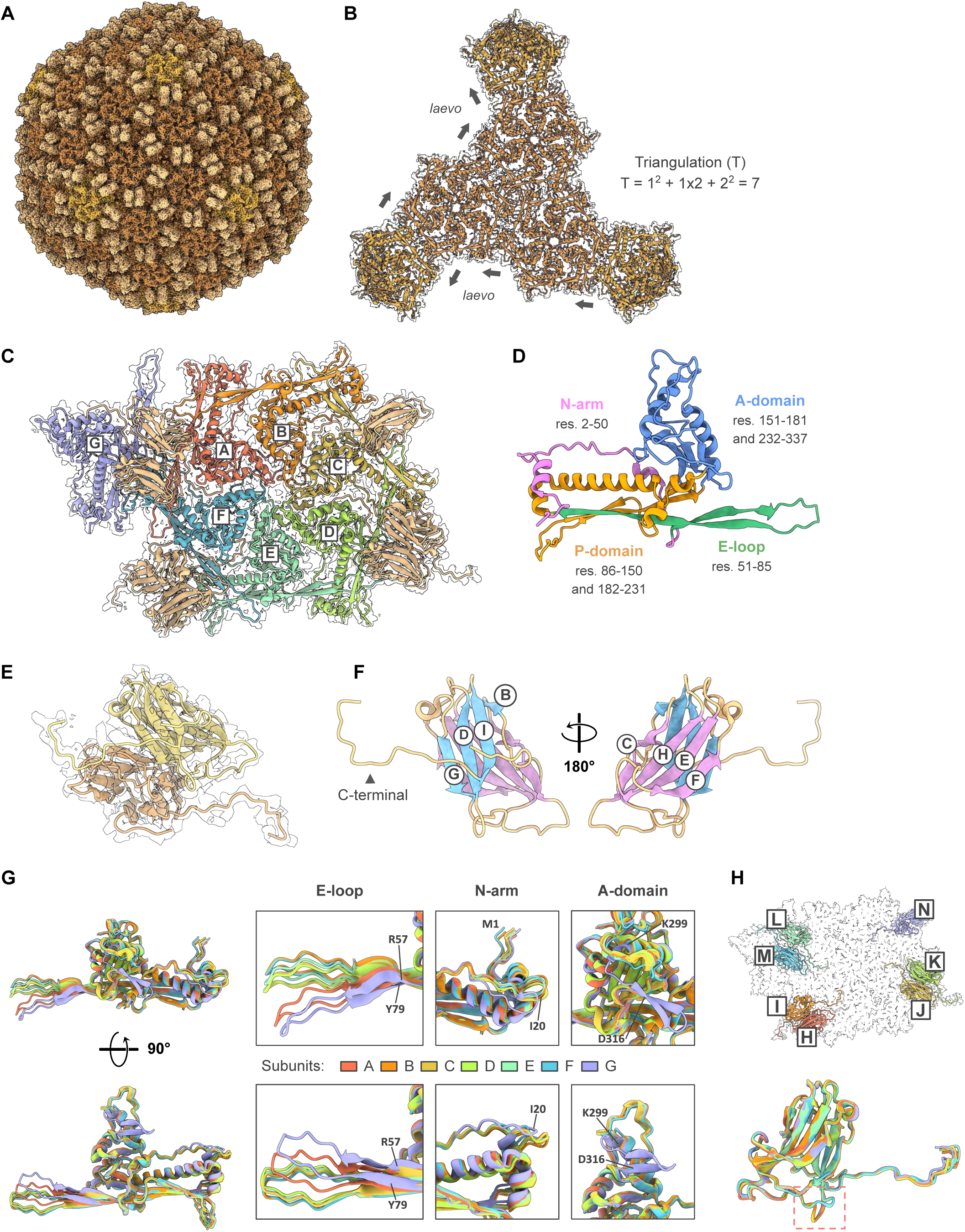
Capsid architecture of ΦXacm4-11. **(A)** Surface map of the capsid assembled with icosahedral symmetry (I), viewed along a 3-fold axis. Capsomers are differentially coloured to distinguish hexons (orange), pentons (yellow-orange), and the cement protein (beige). **(B)** Capsid facet fitted into the cryo-EM density depicting the triangulation number (*T* = 7), the *laevo* lattice hand and the connectivity of capsomers. **(C)** Asymmetric unit (ASU) of the capsid, comprising seven MCP subunits, six contributed by an hexon and one by an adjacent penton, with seven copies of the cement protein, shown fitted into the cryo-EM map. **(D)** Domain organization of the HK97-like MCP, highlighting the N-terminal arm (N-arm), axial (A-) domain, proximal (P-) domain, and E-loop, with residue ranges indicated. **(E)** CP dimer fitted into the cryo-EM density. **(F)** Orthogonal views of the CP dimer, with individual β-strands labelled to illustrate molecule topology. **(G)** Superposition of the seven copies of the MCP within the ASU. Three major locations are highlighted: the E-loops (residues Arg57-Tyr79, left), the N-arm (residues Met1-Ile20, center) and the annular loop (residues Lys299-Asp316, right). **(H)** *Top:* Contribution of seven CP to the ASU of the capsid as in **(C)**, shown fitted into the cryo-EM map. *Bottom:* Superposition of the seven copies of the CP within the ASU, highlighting the only non-overlapping flexible turn. RMSD values can be found in **Table 2**.

**Table 2.** Comparative RMSD values (all atoms) for major capsid proteins (subunits A-G) and for cement proteins (subunits H-N) of the asymmetric unit (ASU).

|  | **A** | **B** | **C** | **D** | **E** | **F** | **G** |
| --- | --- | --- | --- | --- | --- | --- | --- |
| **A** | - | 2.879 | 2.201 | 2.031 | 2.285 | 2.847 | 4.148 |
| **B** |  | - | 1.584 | 1.614 | 1.387 | 1.233 | 5.523 |
| **C** |  |  | - | 1.203 | 1.343 | 1.598 | 4.944 |
| **D** |  |  |  | - | 1.286 | 1.756 | 4.949 |
| **E** |  |  |  |  | - | 1.500 | 5.038 |
| **F** |  |  |  |  |  | - | 5.190 |
| **G** |  |  |  |  |  |  | - |

|  | **H** | **I** | **J** | **K** | **L** | **M** | **N** |
| --- | --- | --- | --- | --- | --- | --- | --- |
| **H** | - | 0.836 | 0.895 | 0.884 | 1.433 | 1.463 | 1.008 |
| **I** |  | - | 0.957 | 0.814 | 1.446 | 1.458 | 0.931 |
| **J** |  |  | - | 0.805 | 1.430 | 1.421 | 0.873 |
| **K** |  |  |  | - | 1.443 | 1.454 | 0.932 |
| **L** |  |  |  |  | - | 0.878 | 1.513 |
| **M** |  |  |  |  |  | - | 1.532 |
| **N** |  |  |  |  |  |  | - |

A representative model of the full-length MCP and CP (residues 2-337 and 1-148, respectively) illustrates their structural features (**Figure 3D,F**). The ΦXacm4-11 MCP has an HK97-like fold that is observed in the capsids of many other bacteriophages [44]. This topology has the following features: N-terminal arm (N-arm), extended loop (E-loop), P-domain and A-domain (**Figure 3D**). The CP has a canonical jelly-roll fold topology made up of eight beta strands (B-I) arranged in two four-stranded antiparallel sheets (β strands BIDG and CHEF) with an 18 residue C-terminal extension (**Figure 3F**).

Superposition of the seven copies of the major capsid proteins showed they are also very similar except for three regions (**Figure 3G** and **Table 2**). The MCP E-loops, corresponding to residues Gly51-Ser85, are very similar in subunits B-F while in subunits A and G the loop is bent downward as it emerges from the main globular portion of the MCP. This could be due to the fact that these two subunits are related by a pseudo-two-fold axis at the interface between the pseudo-six-fold capsomere (hexon) and the five-fold capsomere (penton), while all other MCP subunits are found at the interfaces between two pseudo-six-fold capsomeres. Another related difference between MCPs can be found at the N-arm (residues Met1-Thr50) that interacts directly with the loops at the base of the CP jelly roll fold. Again, the greatest difference in this N-terminal region is found in between subunit G found in the pentons and subunits A-F found in the hexons. A third major difference between MCPs is the conformation of the annular loop corresponding to residues Lys299-Asp316 in the C-terminal region of the protein (axial-domain) (**Figure 3G**). This loop adopts an open configuration in subunits A-F while in subunit G it forms a more compact, hydrogen-bonded hairpin. Again, the difference is explained by the fact that this loop is located at the 5-fold vertex made up of 5 equivalent MCP G subunits at the centre of the pentons and at the pseudo 6-fold vertex at the centre of the hexons made up of MCP A-F subunits (**Table S3**). Superposition of the seven copies of the CP in the asymmetric unit shows that they are all very similar, with RMSD between 0.8 Å and 1.5 Å (**Figure 3H** and **Table 2**). The largest RMSDs are found between subunits L and M, located at the edges between pentons and hexons, and subunits H, I, J, K and N, that are found at the edges between two hexons (**Table S3**).

### Cryo-EM structure of ΦXacm4-11 portal-tail complex

In order to obtain more information regarding the composition and molecular structure of the phage tail complex, we carried out another single particle reconstruction workflow using particles previously subtracted with a mask localised in this region (**Figure S3**), resulting in a 3.45-Å resolution map that allowed us to identify and model several of its constituent structural proteins (**Figure 4** and **Figure S5B,C**).

**Figure 4.**
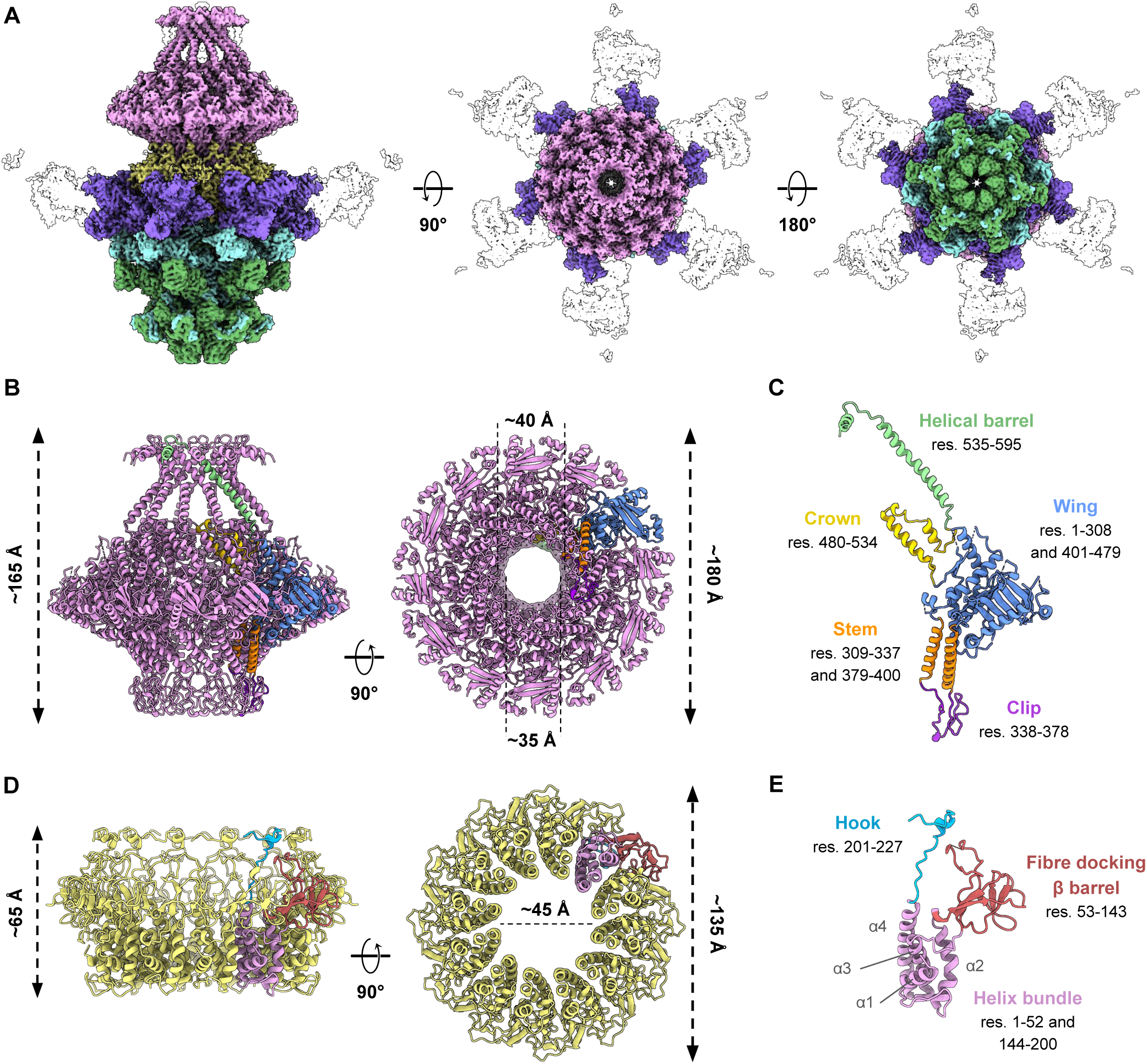
Structural organization of the tail-portal region of ΦXacm4-11. **(A)** Cryo-EM reconstruction of the phage tail, displayed as a density map coloured according to the fitted gps. *Left to right*: side, top and bottom views. **(B)** Atomic model of the portal complex assembled with C12 symmetry. *Left to right*: side and bottom views. **(C)** Portal protein monomer (gp22), coloured by its structural domains. **(D)** Atomic model of the adaptor assembly in C12 symmetry. *Left to right*: side and bottom views. **(E)** Adaptor protein monomer (gp29), coloured by its structural domains. Individual domains highlighted in (B) and (D) are colour-coded as in panel (C) and (E), respectively.

The portal-tail complex consists of, at least, five different proteins organised in sections ordered starting from the inside to the outside of the capsid: the portal (gp22), the adaptor (gp29) plus the tail fibres (gp36), and the distal tip of the tail called the nozzle composed of gp30 and gp31 (**Figure 2** and **Figure 4A**).

### Portal

The portal protein gp22 forms a dodecameric ring in which the extended monomers are oriented parallel with the channel axis (**Figure 4B**). Each portal subunit can be divided into wing, stem, clip, crown and helical barrel domains (**Figure 4C**) [45]. In our structure, 515 of the 598 residues of each gp22 subunit could be confidently modelled within the EM map. The wing, stem, crown and helical barrel domains are located within the capsid and the central clip domain is embedded within the capsid layer of the special vertex. In addition to N- and C-terminal residues 1-14 and 595-598, unmodeled segments are located in loops in the wing domain. The most significant unmodelled is a loop immediately following the stem (residues 406-425) that projects into the lumen and has been observed to introduce a constriction in the diameter of the channel in other T7-like viruses, thus being termed the “valve” [46,47]. Density corresponding to this unmodelled valve however is observed as an expanded ring around the DNA or DNA-protein complex running down lumen of the channel (see below). Three other constrictions are clearly observed in the portal channel: a ∼28 Å constriction at the top of the channel formed by hydrophilic residues from the C-terminal helical barrel, similar to the “Valve S” found in the cyanophage P-SCSP1u [46], a 37 Å constriction in the centre formed by the crown helix hairpin and another 40 Å constriction at the bottom of the stem. Interestingly, in other T7-like viruses the clip domain further constricts the diameter of the portal channel [46,48]. However, in ΦXacm4-11, the clip domain is splayed outwards to interact with the inner lumen of the adaptor protein (see below), thus increasing the channel diameter to ∼50 Å. These clip domains, each made up of 3 antiparallel beta strands, interact with each other through strand-swapping interactions to form a continuous zig-zag network of hydrogen-bonded strands.

### Adaptor

The adaptor is a dodecameric ring of equivalent subunits coded by gp22 (**Figure 4D**). Each subunit can be divided into three domains, a lower domain made up of 4 α-helices, the first two of which (α1 and α2) are derived from the N-terminal of the polypeptide chain (residues 1-52) while the other two (α3 and α4) come from the C-terminus (residues 144-200). The upper domain consists of a beta sandwich of two antiparallel beta sheets derived from residues 53-143 plus a coiled region, called the hook, containing a small alpha helix derived from the extreme C-terminus (residues 201-227) (**Figure 4E**). Where the portal dodecamer inserts into the adaptor, the former’s clip domains make intimate contacts with the ends of two α4 helices and C-terminal tails from neighboring adapter lower domains as well as with the linker between the lower and upper domains from one adaptor subunit (**Figure 4A,E**). The C-terminal tails extend upwards making more contacts with the portal stem domain helices. The bottom (most distal) point of the adaptor channel is the most narrow (45 Å in diameter) where the parallel α4 helices of the dodecamer contribute to the inner lining of the channel [46,47].

### Interaction of the portal-adaptor complex with the capsid

The twelve-fold portal-adaptor complex is inserted within the five-fold special vertex of the viral capsid (**Figure S4A**). Due to symmetry mismatch between the interacting structures, the features of the tail in the C1 EM map of the whole viral particle are blurred into low resolution rings into which the molecular model of the tail could be inserted, allowing us to detail the interaction of adaptor and portal proteins with the five hexons surrounding the special vertex. While all other hexon-penton interfaces in the capsid icosahedron include a pair of cement proteins (L and M subunits), no density corresponding to the five pairs of cement protein was identified at the special vertex as they would suffer severe steric clashes with the portal-adaptor complex. The interface involves 15 MCP subunits, three from each hexon. One MCP subunit interacts via its P-domain (subunit A), the other with its A-domain (subunit F) and the third contributes with the tip of its E-loop (subunit B). These MCPs form a ring inside which is inserted into a complementary groove on its outer surface of the portal-adaptor complex. This groove’s upper surface is made up of the wing domains of the portal protein that interact principally with the inner lumen of the capsid special vertex. The lower surface of the groove is made up of the upper domain and C-terminal extension of the adaptor subunit that interacts with the outer surface of the capsid (**Figure S4E** and **Movie S1**).

### Interactions of the portal-adaptor complex with DNA

The lumen of the portal-adaptor complex is often called the DNA-ejection channel [49]. In the C1 EM map, we observe a clear columnar or needle-like density within this channel that is ∼32 nm in length, beginning above the core and running down to the adaptor (**Figure S4A**). Along most of its length, the needle has a diameter of approximately 19 Å but increases significantly at both ends: to approximately 68 Å above the core and to 45 Å at the adaptor-nozzle interface. The density also increases in diameter within the portal at the position corresponding to the unmodelled loop that forms the valve construction in other T7-like viruses [46]. This needle-like density has been observed in many other podovirus structures and has been proposed to be due to DNA [50,51] and/or protein (**Figure S4D**). We are not able at this time to identify the chemical nature or identity responsible for this density but its diameter in the narrow portions is consistent with double-helical DNA [52,53]. Interestingly, the expanded density at the tip of the needle in the portal/adaptor complex exhibits an arrowhead-like, fivefold-symmetric (C5) morphology in the C1 map (**Figure S4D**), consistent with a structure specialized for membrane penetration.

### Nozzle (tail tip) complex

The proteins gp30 and gp31 interact to form an intricate unified nozzle structure. Six of these heterodimers come together to produce the 6-fold symmetric end of the tail (**Figures 2, 4, 5** and **6**). Each gp30:gp31 heterodimer presents the same domain composition and topology as previously observed for other podoviridae phage nozzle proteins [47], i.e. platform, β-propeller, fibre dock and tip (**Figure 5F** and **Movie 1**). However, four extra globular domains (EDs) can be found in the ΦXacm4-11 gp30:gp31 nozzle that protrude radially from the main conical part of the structure. Although there is no apparent interaction between these extended domains, overall they create a left-handed helical screw that decorates the ΦXacm4-11 tail. The two uppermost domains, ED1 and ED2, each form a β-sandwich composed of seven β-strands: ED1 is derived from gp31 (residues Gly307-Arg406), whereas ED2 is formed by gp30 (residues Thr96-Val175). The remaining two domains, ED3 and ED4, also adopt β-sandwich folds but are composed of contributions from both nozzle subunits.

**Figure 5.**
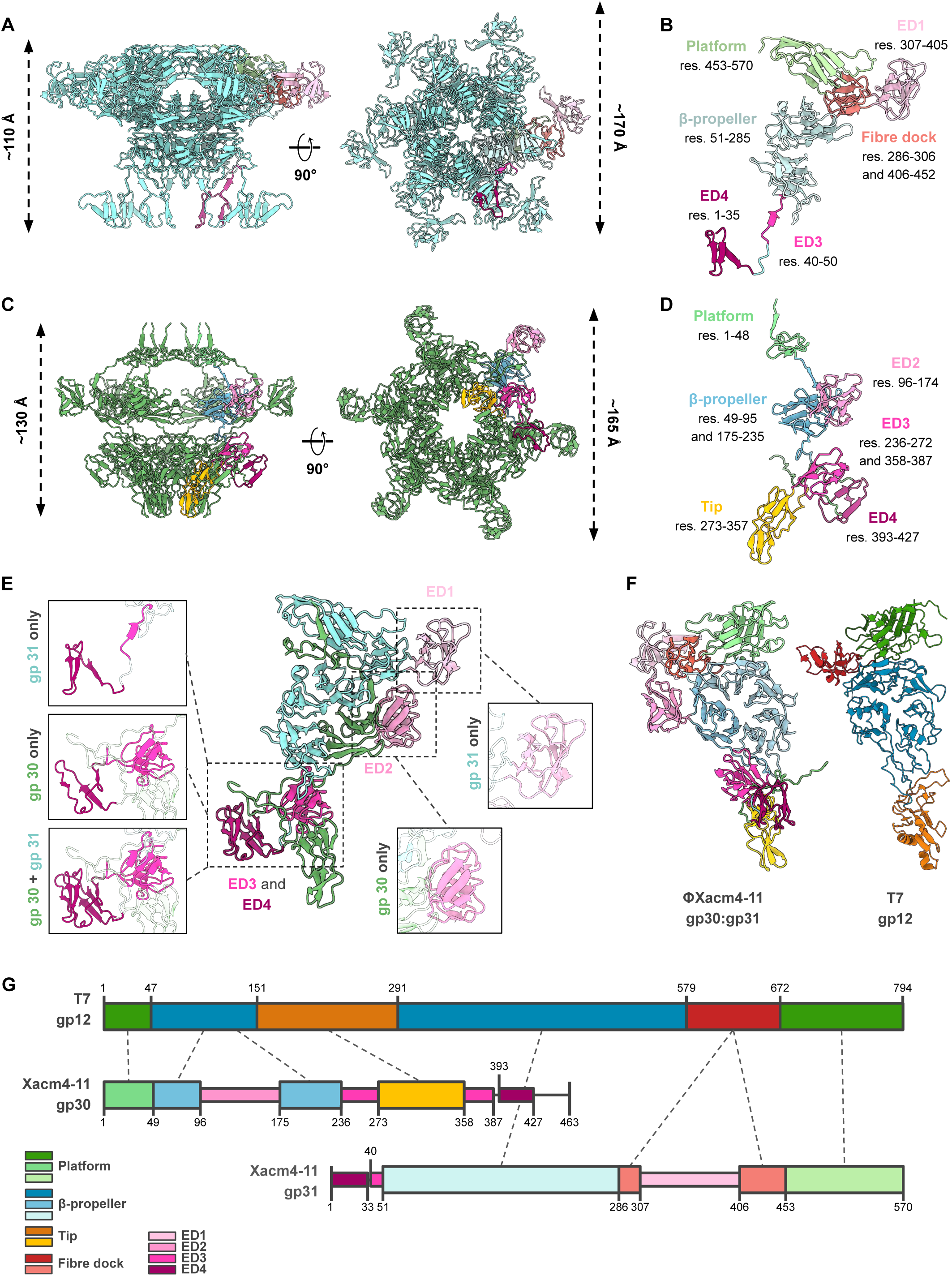
Components of the ΦXacm4-11 nozzle complex, a hexamer of gp30-gp31 dimers. **(A)** Atomic model of the nozzle subunit gp31 complex assembled with C6 symmetry. *Left to right*: side and bottom views. **(B)** Monomer of gp31 coloured by structural domains. **(C)** Atomic model of the nozzle subunit gp30 assembled with C6 symmetry. *Left to right*: side and bottom views. **(D)** Monomer of gp30, coloured by its structural domains. Individual domains highlighted in (B) and (D) are colour-coded as described in panel (A) and (C), respectively. **(E)** Atomic model of the gp30-gp31 heterodimer, highlighting the formation of extension domains (EDs) at the interface. ED1 is formed exclusively by the gp31 and ED2 by gp30, whereas ED3 and ED4 have contributions from both subunits. **(F)** Structural comparison between the ΦXacm4-11 gp30-gp31 and the homologous single gp12 subunit from bacteriophage T7. **(G)** Schematic comparison of domain organization between T7 gp12 and ΦXacm4-11 gp30 and gp31. Coloured blocks indicate corresponding structural domains and extension regions, with dashed lines marking inferred homologies and domain rearrangements between systems. All panels use a consistent colour scheme across maps, atomic models and schematics to facilitate comparison.

ED3 is formed primarily by gp30, comprising six β-strands (residues Asp236-Lys273 and Thr359-Val387) complemented by a single β-strand contributed by gp31 (residues Thr40-Arg51). In contrast, ED4 is assembled from interwoven β-strands derived from both proteins, with three β-strands from gp30 (residues Pro393-Gly427) and three β-strands from gp31 (residues Ala2-Ala33) forming the β-sandwich (**Figure 5**).

Analyzing the lumen of the tip, we note two gates that secure the content of the virion until release (**Figure 6A**). The lower gate is made up of residues Gly134-Val140 (GGAQDVV) and the upper gate is made up of residues Gln183-Gly188 (QFEKDG), both from gp31. Interestingly, these two gates present different chemical natures for closure. The inner aperture of the upper gate is formed by the hydrophobic interactions between residues Phe184 from the 6 copies of the protein while the lumen of the lower gate lined by the six pairs of side-chains from Gln137, resulting in a more tightly fitted pseudo-3-fold structure (**Figure 6B**). The cavity between these gates appears to be empty, in spite of its significant surface ∼1,500 Å^2^ and volume of ∼3,750 Å^3^. Two other constrictions are observed at the top of the nozzle. One is found immediately below the adaptor, made up of residues Asn464-Met471 (NSPGGDSM) from gp31. This constriction is relatively wider than the previous ones, yet it appears sufficient to prevent the premature release of the DNA-ejection complex (**Figure 6A,B**). The other constriction is a ring of vertical beta hairpins (residues 337-348 of gp30) that form the distal tip of the nozzle (**Figure 5C,D** and **Figure 6A,B**).

**Figure 6.**
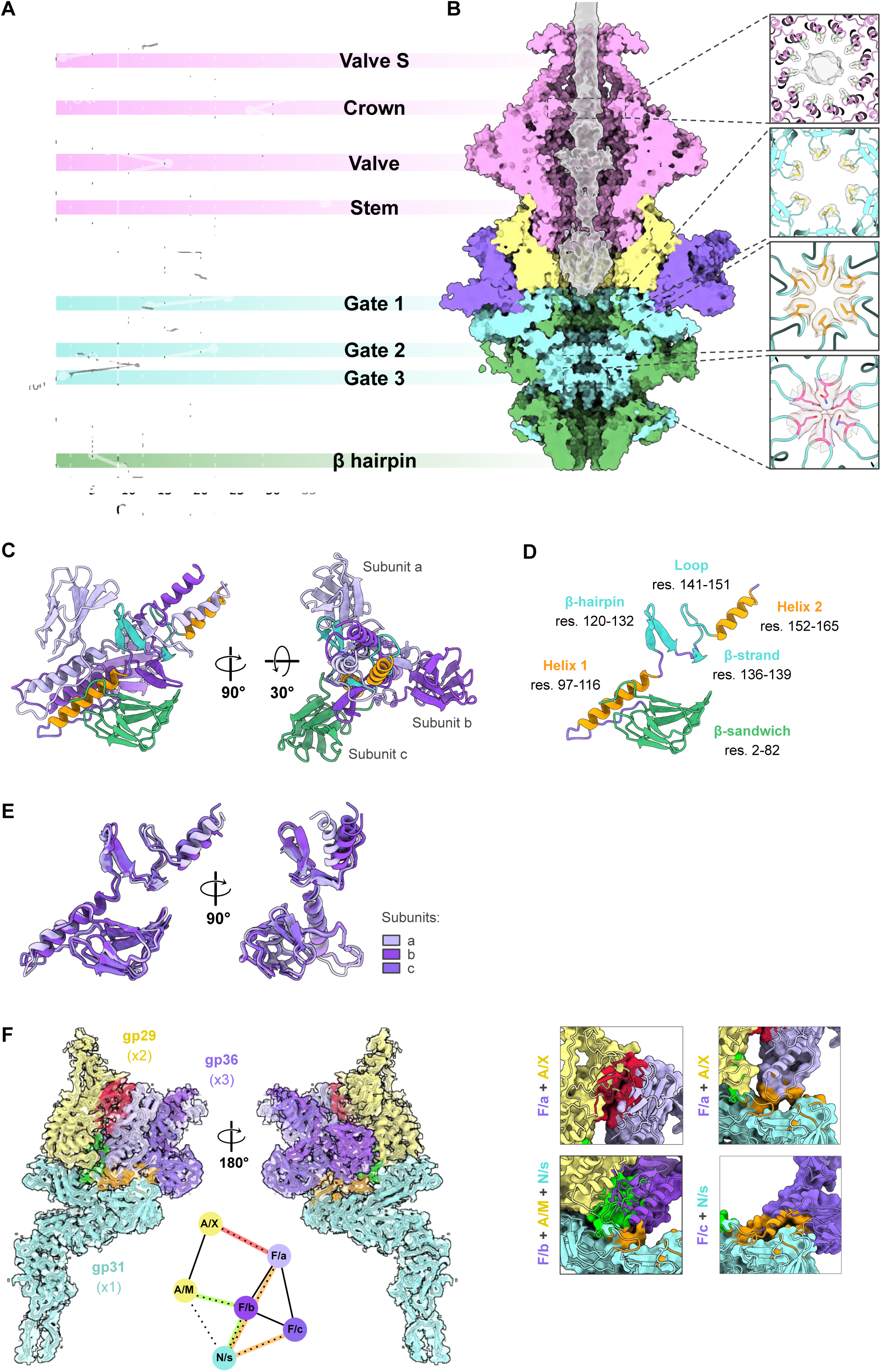
Internal channel architecture of the ΦXacm4-11 tail and organization of the proximal fiber. **(A)** Channel radius along the phage tail axis (z). The narrowing points within the internal conduit determine distinct structural regions. Three main constrictions at the lower third of the tail were characterized as gates 1, 2 and 3. **(B)** Longitudinally-sectioned surface representation of the tail cryo-EM map, with cross-sections illustrating the chemical composition of the inner channel at selected positions. **(C)** Atomic model of the trimeric N-terminal region of the proximal fiber protein (gp36), shown in two lateral orientations. **(D)** Architecture of the N-terminal region of the gp36 monomer coloured by domain. **(E)** Superposition of the N-terminal regions of the three subunits within the gp36 trimer, showing high structural similarity with minor divergence at helix 2. **(F)** Model of the interaction between two adaptor subunits, one fiber trimer and one nozzle protein subunit gp31, fitted into the cryo-EM density, with interface surfaces highlighted. An interface diagram indicates distinct adaptor and nozzle contacts for each fiber subunit. The nozzle gp30 subunit does not participate directly in interactions with the tail fibre gp36 trimer and is not shown for clarity.

We note that the C-terminal of gp30 (e.g. Pro432) is very close (∼6 Å) to the N-terminal of gp31 (e.g. Gly4), indicative of a circularly permuted polypeptide. This is supported by the fact that the last 32 residues of gp30 (PNPYYGGGGGVNGSGGGSVRPGGSSNVQQEAL) is highly rich in glycine and serine residues (46.9% in total) coded by repeatedly used codons (GGC 91% and GGG 9% for glycine and AGC 100% for threonine). This is consistent with the observation that in several phages, such as the archetypal T7 phage, the tail tip and nozzle proteins are fused into a single polypeptide chain (gp12; **Figure 5F** and **Movie 1**). This fusion is also observed in several prophages identified via database searches in a wide variety of α-, β-, and γ-proteobacteria species, such as *Pseudomonas hormoni*, *Pseudomonas putida* KT2440, *Neorhizobium galegae* bv. *orientalis* HAMBI 540, *Pyruvatibacter mobilis*, *Megasphaera hexanoica*, *Azorhizobium caulinodans*, *Oxalobacter vibrioformis* and *Agrobacterium tumefaciens* S33.

### Tail fibers

The annotation of the ΦXacm4-11 genome includes four predicted tail fibre proteins, gp36-gp39, all of which were confidently detected in the proteomic analysis of purified page particles (**Table 1** and **Table S1**). Inserted in between the adaptor and the nozzle, are six trimers of the tail fibre protein gp36 whose N-terminus is a homolog of the N-terminus of the bacteriophage T7 tail fiber protein gp17 (pfam03906; protein size 553 residues; [50,54]). While these ΦXacm4-11 gp36 are 369 residues in length, we were only able to model the polypeptide chains of their N-terminal halves (residues 2-166) with confidence in the cryo-EM map at 3.5 Å resolution. The N-terminal halves of each gp36 subunit come together to form a trimeric complex with C3 symmetry. Each subunit begins with an N-terminal β-sandwich (residues 2-82) followed by two helices (residues 97-116 and 152-165) that are collinear and come together to form a two-segmented 3-helix bundle. The helical segments are separated by a three-fold strand-swapping barrel-like structure in which each subunit contributes alternatively a β-hairpin (residues 120-132), a lone β-strand (residues 136-139) and a loop (residues 141-151) (**Figure 6C-D**). Interestingly, the N-termini of the three subunits adopt nearly identical topologies, with very low pairwise RMSD values (1.739-2.343 Å), even though they engage in distinct interaction interfaces, with two subunits contacting both the adaptor gp29 and the nozzle protein gp31, and the third contacting only the latter (**Figure 6E-F**).

In spite of the reduced quality of the EM map beyond residue 166 of gp36, clear density corresponding to molecular components is observed further along the tail fibers (**Figure 2A,C** and **Figure 4A**). To further investigate the composition of the tail of ΦXacm4-11, we generated AlphaFold3 [55] models of the trimeric full-length gp36 protein, superposed them on the experimentally determined models of the N-terminal-domains and adjusted the C-terminal domains of the trimer into the EM map density (**Figure 7A** and **Movie 2**). The C-terminal halves of each gp36 subunit have two distinct β-rich domains: a 6-stranded β-sandwich spanning residues Lys174 to Leu244 followed by a β-helix-like domain extending from Ile264 to Gln368, composed of eleven β-strands. Residues 245-263 are disordered, forming a flexible linker connecting these domains (**Figure 7B**). Within each trimer, structural comparison reveals a high degree of internal symmetry, with RMSD values of 1.051 Å for the β-sandwich domains and 0.903 Å for the β-helix-like domains (**Figure 7C**). This arrangement gives rise to two trefoil-like architectures: one formed by the three β-sandwich domains and another by the three β-helix-like domains, each contributed by one gp36 subunit (**Figure 7D**). The combination of this intricate domain topology and the conformational flexibility provided by the inter-domain linkers enables the gp36 trimer to adopt a ∼60° kink observed in the cryo-EM map of the tail fiber (**Figure 7E**).

**Figure 7.**
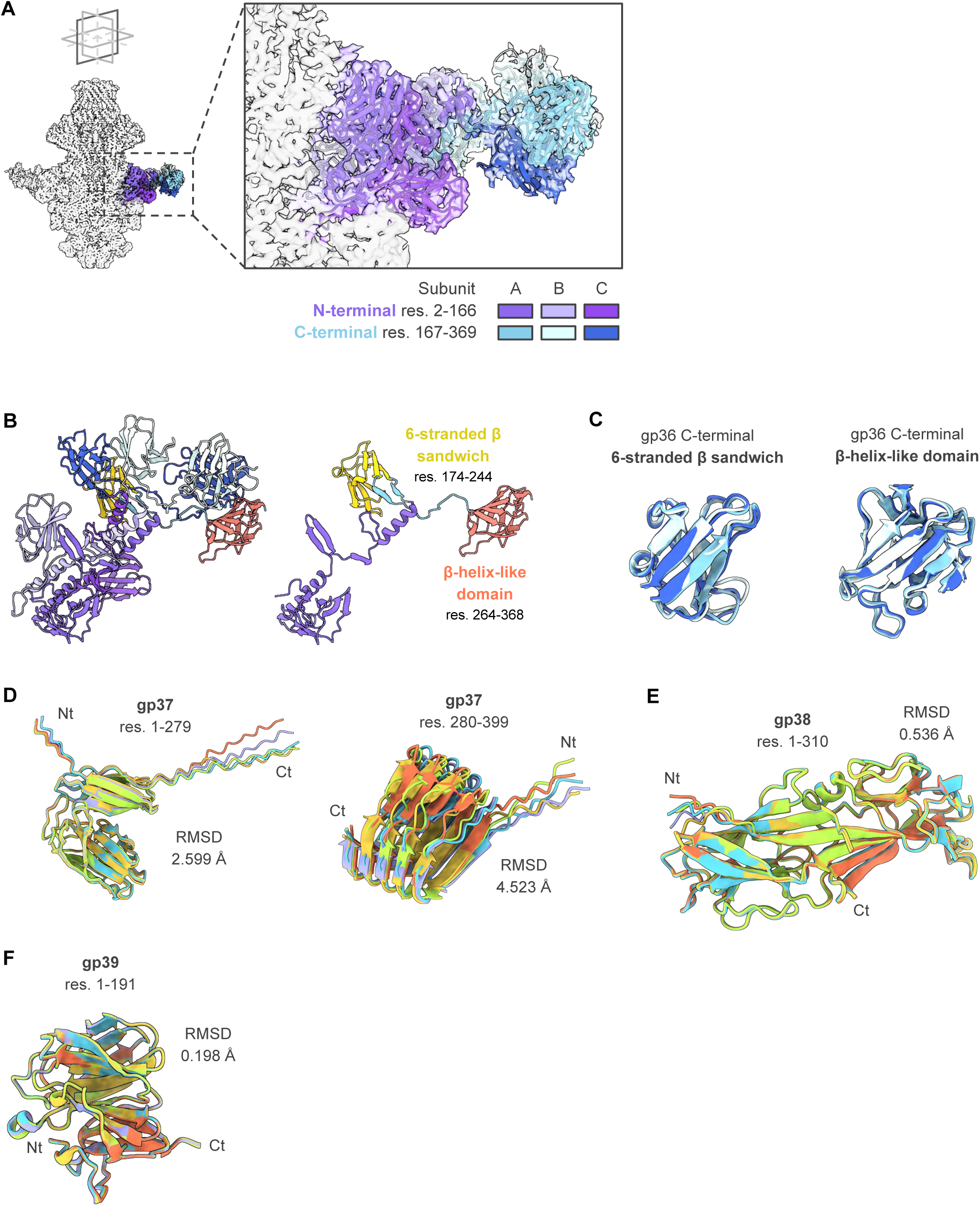
Proximal fiber architecture and predicted fibre components of ΦXacm4-11. **(A)** Cryo-EM map of the tail with one full-length gp36 trimer fitted into the density. **(B)** *Left to right*: Full-length gp36 shown as a trimer and as a monomer, with the C-terminal domains indicated. **(C)** *Left to right*: Superposition of the β-sandwich and the β-helix-like domains, both from the C-terminal of gp36, revealing high structural similarity. The domains were treated independently due to flexible inter-domain linkers, which result in distinct relative orientations among subunits within the trimer. **(D-F)** AlphaFold3-predicted structures of gp37 (D; N- and C-terminal regions shown separately), gp38 (E), and gp39 (F). For each protein, the five predicted structures are shown superimposed in a 3D alignment. RMSD values are indicated in each case.

The remaining density observed in the tail fibers EM map is most plausibly attributed to the tail fiber proteins gp37, gp38, and gp39. However, these three proteins have no known homologs identifiable by sequence similarity. We therefore used their AlphaFold-generated models to query the pdb structural database using the Dali server [56] (**Figure 7F-H**). This procedure identified potential carbohydrate binding modules or folds encountered in phage tails in all three proteins which we now describe briefly. Gp37 is predicted to have two N-terminal β-sandwich domains (**Figure 7F**), one of which has a topology equivalent to known oligosaccharide binding modules, for example those identified in a mannanase from the thermophile *Thermoanaerobacterium polysaccharolyticum* (PDB: 2zew, [57]) and *Thermotoga maritima* endo-β-1,4-galactanase (PDB: 2xom, [58]), including acidic residues involved in Ca^2+^-binding. The C-terminal domain of gp37 is predicted to adopt a β-helix of five rungs with two strands in each rung (**Figure 7F**). Phage related proteins with carbohydrate binding domains and extended β helices have been observed, for example the bacteriophage CBA120 tailspike protein 4 (PDB: 5w6h, [59]), the *Klebsiella* phage Kp7 type II tail fiber gp52 (PDB: 7xyc, unpublished) and the *Klebsiella pneumoniae* phage K2-2 tailspike depolymerase (PDB: 8iqe, [60]). The predicted structure of gp38 (**Figure 7G**) has only partial topological similarity with the baseplate component gp7 from phage T4 (PDB: 5hx2, [61]), the protein fiber I from *Ralstonia solanacearum* phage GP4 (PDB: 8jou, [62]), and te whisker protein from *Shigella flexneri* phage Sf14 (PDB: 9c39, [63]). Finally, gp39 is predicted to be a 3-bladed β-propeller (**Figure 7H**) and has a similarity with 7-bladed β-propeller lectins (PDB: 8q82, [64]; PDB: 7c3c, [65]; PDB: 4agi, [66]) and sialidase enzymes from *Bacteroides acidifaciens* (PDB: 7mhu, [67]) and *Ruminococcus gnavus* (PDB: 4×47, [68]). We attempted to insert the models of these three tail fiber proteins (gp37, gp38 and gp39) into the remaining density that extends beyond the C-termini of the gp36 trimer. However, these efforts were inconclusive due to map resolution and uncertainties regarding the stoichiometry and specific protein-protein interaction interfaces. Definitive molecular models of the distal regions of the ΦXacm4-11 tail fibers will require further experimental investigation.

### Candidates for ΦXacm4-11 core proteins

Upon contact with the target cell, the double-stranded phage DNA will pass through the lumen of the portal tail complex. In mature phage particles, ejection is restricted by constrictions that reduce the diameter and hold in place proteins that plug the channel [47,69,70]. Furthermore, the tails of podophages are too short to cross the bacterial cell envelope and access the cytosol; e.g. for ΦXacm4-11 the tail is ∼20 nm in length. To overcome this limitation, proteins found inside the viral capsid (core or pilot proteins) disassemble and are injected into the bacterial membrane and periplasm where they reassemble to extend the DNA translocation channel [71,72]. In phage T7, these core proteins are gp14 (196 residues), gp15 (747 residues) and gp16 (1,317 residues) and have been proposed to exist in a 8:8:4 complex sitting atop the portal in a dome-like structure within the mature capsid [50,51] but can form an alternative-tube like configuration *in vitro,* interpreted to correspond to that adopted upon translocation into the target cell [71–73].

In the C1 map, density within the capsid above the portal protein can be detected which we assume corresponds to the ΦXacm4-11 core proteins. In T7, the genes coding for these three proteins are found between those for the nozzle and filament proteins gp12 and gp17, respectively. In ΦXacm4-11, these positions correspond to ORFs 33, 34 and 35, which remain syntenically conserved. While ΦXacm4-11 gp33 (167 residues) and gp34 (730 residues) are similar in size to their T7 analogs gp14 and gp15 respectively, ΦXacm4-11 gp35 (2,300 residues) is significantly larger than T7 gp16. Pairwise alignments revealed little sequence conservation between these proteins in T7 and ΦXacm4-11 (**Figure S6**). However, the AlphaFold3 model for gp33 is highly similar to gp14 of T7 phage where it occurs in six copies in an extended helical hairpin conformation as part of the tail complex after DNA ejection [71,73] (**Figure S7**). For gp34, searching its Alphafold3 model against the PDB using DALI gave many hits against several peptidoglycan transglycosylases (Z-scores up to 8.1), including the globular domain of the gp16 subunit of the T7 DNA ejectosome periplasmic tunnel (PDB: 7k5c (Z-score 6.2), [73] and PDB: 6yt5 (Z-score 5.2), [71]). We could not encounter any significant similarities of the Alphafold model for gp35 with proteins in the Protein Data Bank.

## CONCLUDING REMARKS

In this study, we combined genomic annotation, proteomics and high-resolution cryo-electron microscopy to provide the first detailed structural description of a *Xanthomonas*-infecting podovirus. The ΦXacm4-11 virion displays a canonical T7-like architecture while incorporating distinctive features within its portal-tail complex that are likely linked to its infection strategy.

The asymmetric reconstruction revealed a well-organized DNA-protein injection machinery embedded at the unique fivefold vertex of the capsid. In particular, the expanded density observed at the distal end of the portal-adaptor-needle assembly adopts a fivefold-symmetric, arrowhead-like morphology, consistent with a structure specialized for membrane penetration during genome delivery. Such an architecture supports a model in which ΦXacm4-11 compensates for its short tail by deploying an internal injection device to traverse the bacterial cell envelope, analogous to mechanisms proposed for other T7-like phages.

As pointed out by Balogh *et al.* (2013) [16], ΦXacm4-11 is not a typical citriphage since it was not isolated from citrus canker lesions, but rather from *X. alfalfae* subsp. *citrumelonis* strain 36 (Xacm36) lesions. It also differs from most citriphages by its wide host range [16]. While *X. citri* infection by ΦXacm4-11 is type IV pilus-dependent, the molecular basis of this promiscuity is not yet known. As the prevalence of antibiotic-resistant bacterial strains continues to escalate, understanding the dynamics of bacteriophage-host interactions becomes imperative for the development of alternative therapeutic approaches. ΦXacm4-11 is emerging as a good model to study the structure-specificity relationships in T7-like phages.

Together, these findings provide a structural framework for understanding Type IV pilus-dependent infection by podoviruses and highlight ΦXacm4-11 as a powerful model to investigate structure-function relationships in phage entry. This work lays the groundwork for the rational exploitation of *Xanthomonas* phages in biocontrol strategies targeting economically important phytopathogens.

## MATERIALS AND METHODS

### Phage propagation and preparation

Phage ΦXacm4-11 particles were isolated from susceptible propagating bacteria in soft-agar plates. Briefly, an overnight 2xTY (16 g/L tryptone, 10 g/L yeast extract, 5 g/L NaCl) culture of *Xanthomonas citri* strain 306 [36] (∼1×10^9^ CFU/mL) was mixed with ΦXacm4-11 particles (1×10^5^ PFU/mL) at a multiplicity of infection between 0.0001 and 0.00001. After incubation for 10 minutes at 28 °C, 600 µL of the mixture were added to 15 mL of warm KB soft-agar medium (20 g/L peptone, 1.5 g/L K_2_HPO_4_, 15 g/L glycerol, 0.7% agar, pH 7.0) supplemented with 2 mM CaCl_2_ and 6 mM MgSO_4_. The whole mixture was then layered over a previously prepared plate containing 40 mL of KB agar medium (1.5% agar) supplemented with 2 mM CaCl_2_, 6 mM MgSO_4_ and 100 µg/mL ampicillin, and incubated at 28 °C until lysis plates were evident. After this time, phage particles were extracted from the top layer using buffer SM (30 mM Tris, 3% NaCl, 10 mM CaCl_2_, 10 mM MgSO_4_, pH 7.5), and precipitated with 4% PEG 8000 and 500 mM NaCl for 24 hours at 4 °C. Particles were then pelleted by centrifugation for 1 hour at 35,000 xg and 4 °C, and resuspended in buffer SM. A step of CsCl gradient ultracentrifugation for 24 hours at 150,000 g and 4 °C was carried out for further purification. Finally, ΦXacm4-11 phage particles were concentrated to desired titer and buffer exchanged to fresh SM using a 100-kDa Amicon purification system (Merck-Millipore). Sample protein concentration was determined using BCA Protein Assay Kit (71285, Sigma).

### One-step growth assay

The objective of this assay was to determine the burst size, eclipse period and latent period associated with the lytic growth of phage ΦXacm4-11 in *X. citri* strain 306. The one-step assay was performed at 30 °C as described [74] with modifications. Briefly, a phage suspension was added to *X. citri* cultured in 2xTY (16 g/L tryptone, 10 g/L yeast extract, 5 g/L NaCl) at a multiplicity of infection (MOI) of 0.001. After incubating at 30 °C for 10 minutes to allow phage adsorption, the mixture was centrifuged for 1 minute at 6,000 rpm (2,100 xg) at 4 °C. The supernatant was collected for determining the fraction of non-adsorbed phages. The pellet was resuspended in 20 mL of KB medium (20 g/L peptone, 1.5 g/L K_2_HPO_4_, 15 g/L glycerol, pH 7.0) supplemented with 2 mM CaCl_2_, incubated at 30 °C with slow shaking (100 rpm), and 300-μL samples were collected every 10 minutes. These samples were centrifuged at 13,000 xg for 1 minute at 4 °C and the supernatant was diluted in 10-fold series for determination of plaque-forming units (PFU). Results were expressed as PFU/mL at each time point.

### Genome sequencing, assembly, annotation and analysis

Phage genomic DNA was purified from high-titer lysates using silica-based DNA extraction columns, following the manufacturer’s instructions (Thermo Fisher Scientific, Waltham, USA). Purified DNA was quantified and submitted for whole-genome sequencing using a MiSeq Illumina platform (Illumina, San Diego, USA), generating paired-end reads. Raw sequencing data were subjected to quality control and preprocessing, including adapter trimming and removal of low-quality reads, using standard pipelines available within the Galaxy platform [75]. High-quality reads were then assembled *de novo* to generate a single contiguous genome sequence.

Genome annotation was initially performed using automated annotation tools available through the Galaxy framework [75]. Open reading frames (ORFs) were predicted using phage-appropriate gene-calling algorithms. Functional annotation was subsequently refined by extensive manual curation, based on sequence similarity searches at both the nucleotide and protein levels. Putative gene functions were assigned through comparative analyses against publicly available databases, including NCBI (non-redundant nucleotide and protein databases) and KEGG, using BLAST-based approaches [76,77]. Protein domains and conserved motifs were evaluated to support functional predictions. When no significant similarity was detected, ORFs were annotated as hypothetical proteins. This combined automated and manual curation strategy ensured a high-confidence functional annotation of the ΦXacm4-11 phage genome.

In order to confirm whether ΦXacm4-11 genome is linear or circular, we carried out an analysis based on polymerase chain reaction (PCR) amplicons. Briefly, PCR reactions (final volume 25 μL) were performed using 1 μL of ΦXacm4-11 CsCl-purified particles (1×10^8^ PFU/mL) as template (∼50 ng of DNA), 1 μL of in-house produced *Thermus aquaticus* (Taq) DNA Polymerase (1 U/μL), 10 mM Tris-HCl (pH 8.3), 50 mM KCl, 1.5 mM MgCl_2_, 0.001% Gelatin, 0.2 mM dNTPs and 0.3 μM of each oligonucleotide (**Table S4**), in a Mastercycler^®^ Nexus Gsx1 Thermal Cycler (Eppendorf, Hamburg, Germany) under the these conditions: 94 °C for 3 minutes, followed by 30 cycles of 94 °C for 30 seconds, 58 °C for 30 seconds and 72 °C for 5 minutes, and final extension step at 72 °C for 10 minutes. Amplicons were analysed in 1% agarose gels doped with SyBr Safe (Thermo Fisher Scientific) in Tris-acetate-EDTA buffer for 45 min at 80 V.

### Mass spectroscopy analysis

Proteins present in purified phage particles were identified following two separate strategies, as described previously, [78,79]. For the analysis of total proteins in solution, ∼50 µg of purified phage particles (2.4×10^12^ PFU/mL) were precipitated and resuspended in 20 µL of 8 M urea. The sample was reduced and alkylated using 50 mM dithiothreitol (DTT) and 50 mM iodoacetamide (IAA), adjusted to a buffer consisting of 1.5 M urea, 25 mM Tris, 25 mM MgCl_2_, 50 mM NaCl, pH 8.0, and subjected to digestion with trypsin (final trypsin to protein ratio of 1:50) for 16 hours at 37 °C. The reaction was stopped by adding 10% formic acid, desalted using ZipTip C18 (ZTC18S096, Millipore) and finally reduced to dryness by vacuum centrifugation until use. For the analysis of SDS-PAGE resolved bands, in-gel digestion was carried out. Purified phage particles (∼10 µg) were subjected to separation in 15% Tris-Tricine SDS-PAGE [80], fixed and stained using Coomassie Blue G-250. The lane was divided into seven sections according to band sizes. Excised gel pieces were transferred to clean tubes and iteratively washed using 200 µL of 50 mM NH_4_HCO_3_ in 40% acetonitrile (ACN) until complete destaining. After a last wash using 200 µL 100% ACN and removal of the solvent, the gel pieces were treated with 10 mM DTT and 100 mM IAA for 30 minutes at room temperature in the dark. Reduced and alkylated gel pieces were then washed twice with rounds of 200 µL 100% ACN and 200 µL 50 mM NH_4_HCO_3_, and treated for 16 hours at 37 °C with 100 ng trypsin in buffer 10 mM NH_4_HCO_3_, 5% ACN. The reaction was then stopped by adding 10% formic acid, and the trypsinized bands were extracted from the gel using several rounds of buffer 40% ACN, 0.1% formic acid, pooled, desalted using ZipTip C18 (ZTC18S096, Millipore) and finally reduced to dryness by vacuum centrifugation until use. In all cases, the final dried samples were resuspended in 10 µL of 5% ACN, 5% formic acid.

Liquid chromatography tandem mass spectrometry (LC-MS/MS) analyses were performed on a Linear Trap Quadrupole (LTQ) Orbitrap Velos ETD (Thermo) coupled with an Easy-nLC II (Thermo) instrument located at Centro de Facilidades de Apoio a Pesquisa of Universidade de São Paulo (CEFAP-USP). The peptides were separated on a C18 Reverse Phase column (Thermo) on a 95-minute gradient. Spectra were extracted and searched using Thermo Proteome Discoverer v1.4.0.288 (Thermo) against both phage (DNA translated) and host protein databases. Software auto validation parameters were used to select confidently identified proteins and peptides.

### Electron microscopy sample preparation and data collection

Negative staining electron microscopy was used to evaluate the quality of the ΦXacm4-11 particles after CsCl purification. Briefly, 3-μL drops of the preparation were deposited onto glow-discharged ultrathin carbon-coated copper grids (Ted Pella, #18024). After incubating the sample for 1 min at room temperature, the grids were rapidly washed three times with H_2_O_dd_, then exposed to a solution of 2% uranyl acetate (twice for 30 seconds each), and stored at room temperature until visualisation and analysis. Images were recorded on either a JEOL JEM 2100 microscope (operating at 200 kV) using a LaB_6_ filament and coupled to a CCD (charge-coupled device) camera.

For cryo-EM sample preparation, 2.5-μL drops of ΦXacm4-11 particles after CsCl purification at 1×10^13^ PFU/mL were applied onto glow-discharged Quantifoil R2/2 200 mesh copper grids (TedPella), and plunge-frozen using Vitrobot Mark IV (Thermo Fisher Scientific) at 4 °C, 100% humidity, and blot time of 4 s. Three independent datasets were collected at the Brazilian Nanotechnology National Laboratory (LNNano) Electron Microscopy Facility on a 300 kV Cs-corrected Titan Krios G3i (Thermo Fisher Scientific). Datasets of 18,031 movies in total were automatically acquired using EPU software (TFS), recorded on a Falcon 3EC direct electron detector (TFS) operated in counting mode with a nominal pixel size of 1.1075 Å/px. The total accumulated dose was of 30 e^-^/Å^2^, divided into either 20 or 60 frames per movie. The defocus range used for all data collections was between-1.0 and-3.0 μm.

### Cryo-EM data processing, refinement and reconstructions

Data processing and reconstructions were performed within the RELION 3.1 software framework [41,81]. Movie frames were aligned and averaged with dose weighting using MotionCor2 [82] to compensate for specimen drift and mitigate the effects of electron beam-induced specimen damage. The contrast transfer function parameters of the aligned micrographs were determined using CTFFIND4.1 [83]. The micrographs were filtered screening for poor Thon rings or resolution estimation worse than 5 Å. Approximately 500 phage particles were manually picked from the selected micrographs to generate representative 2D-class averages, which were used as templates for automated particle picking for the entire data set.

For the icosahedral reconstruction of ΦXacm4-11, a total of 91,832 phage particles were auto-picked and extracted using a box size of 1,000 by 1,000 px, which were downsampled to 850 by 850 px and pixel size to 1.3029 Å/px due to computing limitations. Several rounds of reference-free 2D classification were performed, allowing to keep 71,682 particles that were then subjected to 3D classification generating one good map with ∼97% of them (69,590 particles). This 3D map was then low-pass filtered to 60 Å and used as reference for 3D auto-refinement imposing icosahedral symmetry (I). In order to improve the map quality, anisotropic magnification estimation and correction, re-estimation of per-particle defocus and particle polishing were carried out on the combined dataset (all jobs implemented in RELION 3.1 [81]). For post processing, a solvent mask and a calculated B-factor of-129.74 Å^2^ was applied, improving the overall resolution of the map to 3.16 Å assessed by the gold standard criterion (FSC-0.143). Local resolution estimations in the map were performed with LocalRes inside RELION. By simple inspection of this cryo-EM, it was possible to determine that the triangulation number [84] for ΦXacm4-11 is T7 (*laevo*).

For the reconstruction of the whole ΦXacm4-11 virion, the previous particles were re-extracted using a box size of 1,150 by 1,150 px and downsampled to 850 by 850 px and pixel size to 1.4984 Å/px. After a few rounds of reference-free 2D and 3D classification, a total of 65,055 particles were kept and subjected to 3D auto-refinement not imposing any symmetry (C1) using the previous icosahedral 3D map low-pass filtered to 60 Å as reference. In order to assist this reconstruction, the reference map was oriented in an alternative 52 setting (with the positive Z-axis pointing at the viewer, one of the 5-fold vertices in the Z-axis and another in the positive XZ-plane) [85]. To enhance map quality, anisotropic magnification estimation and correction, re-estimation of per-particle defocus and particle polishing were carried out on the combined dataset. For post processing, a solvent mask and a calculated B-factor of-100.84 Å^2^ was applied, improving the overall resolution of the map to 4.11 Å assessed by the gold standard criterion (FSC-0.143). This map allowed us to easily identify the location of the tail, oriented through the positive Z-axis.

Finally, for the reconstruction of the ΦXacm4-11 tail, the C1-symmetry reconstructed map was segmented and analysed using Segger and UCSF Chimera [86,87], and created a mask surrounding the density of the tail. This mask was used for a particle subtraction job in RELION using a box size of 410 px and the C1-symmetry reconstructed map. The new set of particles was subjected to reference-free 2D and 3D classification and to 3D auto-refinement imposing 6-fold rotational symmetry (C6) using a previously generated 3D map low-pass filtered to 60 Å as reference. Subsequently, CTF refinement and Bayesian polishing were performed on the refined particle set using standard workflows implemented in RELION [81]. For post processing, a solvent mask and a calculated B-factor of-82.57 Å^2^ was applied, improving the overall resolution of the map to 3.45 Å assessed by the gold standard criterion (FSC-0.143). Local resolution estimations in the maps were performed with LocalRes inside RELION.

### Analysis and structure determination

Atomic models of gp25 (major capsid protein), gp26 (cement protein), gp22 (portal protein), gp29 (adaptor protein), and gp30 and gp31 (nozzle proteins) generated by the I-Tasser platform [88] or the ColabFold-AlphaFold2 suite [89,90] were docked and rebuilt into the corresponding cryo-EM map density using UCSF ChimeraX [91,92] and Coot [93] software. For the case of gp36 (tail fibre protein), the incomplete model (residues 2 to 166) was created from scratch by hand using Coot and identifying the amino acid sequence along the cryo-EM map. In all cases, secondary structure predictions were also performed with the PSIPRED server [94] in order to assist the process of model building and identify the protein with a number of secondary structure elements similar to those identified in each cryo-EM map.

For modelling the ΦXacm4-11 capsid, one asymmetric unit (AU), composed of the major capsid protein and the cement protein, was assembled by fitting 7 copies of gp25 (residues 2-337) and 7 copies of gp26 (residues 1-148). Subsequently, the density corresponding to the volume of these 14 molecules within the AU was extracted with UCSF Chimera [87] (using the *zone* tool with a mask radius around the atoms of 5 Å). The atomic models of a single AU were subjected to cycles of refinement in Phenix [94,95] and modelling using Coot until good parameters for Ramachandran plot, MolProbity [96] and EMRinger [97] were obtained. The whole capsid was then created in UCSF Chimera rigidly docking 60 copies of the AU with the Fit-in-Map (*fitmap*) tool, which allowed for the calculation of the tridimensional relationship between AUs using the *measure rotation* tool. This data was subsequently used to calculate the icosahedral symmetry operators and create a BIOMT matrix that permitted the reconstruction of the icosahedral capsid structure for ΦXacm4-11.

A similar strategy was applied to model the ΦXacm4-11 tail. However, the model assembly process was more intricate in this instance, given the varied copy numbers of the proteins constituting the structure. To facilitate the process, the cryo-EM map was initially segmented using Segger inside UCSF Chimera, and saved in three submaps, corresponding to the portal region, the adaptor region, and the rest. In each case, these maps were used to fit and build one subunit of the corresponding proteins. Subsequently, these structures were subjected to cycles of refinement in Phenix and modelling using Coot until good parameters for Ramachandran plot, MolProbity and EMRinger were obtained. Once with all the proteins identified and individually modelled to good validation parameters, the tail structure was created fitting in the cryo-EM map and combining 12 copies of gp22 (residues 15-58, 69-106, 113-206, 215-249, 259-273, 281-406, 426-466 and 471-595), 12 copies of gp29 (residues 2-99 and 110-223), 6 copies of gp30 (residues 2-454), 6 copies of gp31 (residues 2-356 and 363-564), and 18 copies of gp36 (residues 2-166). These protein chains were uniquely named and saved in a simple.pdb file. Finally, new rounds of refinement were carried out in Phenix applying non-crystallographic symmetry (NCS) constraints until good parameters for Ramachandran plot, MolProbity and EMRinger were obtained. A summary of each protein region according to its modeling status is color-coded in **File S3**: transparent indicates unmodelled density, green denotes regions modelled based on cryo-EM data, and yellow corresponds to regions modelled using AlphaFold3 predictions.

### Prediction and in-silico analysis for gp36

The obtained quality for the tail cryo-EM map allowed us to model with sufficient confidence only the amino-terminal region (first ∼165 residues) of the tail fibre protein (gp36). In spite of this, further analysis of the map, using lower contour levels indicates that some extra densities can be found beyond these modelled residues. Due to this, a trimeric structure prediction was carried out using the ColabFold-AlphaFold2 suite, with default parameters. The predicted structure correlates very well with the cryo-EM modelled region, and also displays other two structured regions, all of them connected with two disordered linkers. To broaden our examination, these two regions were isolated using UCSF Chimera, and docked to the extra densities of the map with acceptable correlation. The two linkers were then modelled using Coot, applying local refinement with Torsion, Planar peptide, Trans peptide and Ramachandran restraints. The chimeric version of the tail fibre protein was then subjected to model building into low to medium resolution maps using ISOLDE [98], implemented in UCSF ChimeraX, minimising clashes and other geometric issues.

### Prediction and in-silico analysis of other tail fibre and core predicted proteins

Structure predictions for the tail fibre proteins (gp37, gp38 and gp39) and core proteins (gp33, gp34 and gp35) of bacteriophage ΦXacm4-11 were performed using AlphaFold3 [55] with default parameters. For each protein, five independent models were generated and subsequently evaluated for structural consistency by pairwise superposition and RMSD analysis. Predicted models were visually inspected and analyzed using UCSF Chimera [86,87] and UCSF ChimeraX [91,92]. Structural features such as domain organization, oligomeric state, and surface properties were examined, and regions displaying high confidence and structural conservation among the predicted models were identified. Hydrophobic surface representations were generated to assess the presence of membrane-interacting or channel-facing regions, particularly for core proteins involved in DNA injection. For comparative analyses, representative AlphaFold3 models were structurally aligned to experimentally determined homologous structures available in the Protein Data Bank using UCSF Chimera. Structural similarity was assessed based on overall fold, domain arrangement, and conserved functional motifs. In cases where cryo-EM density was available, predicted models were rigid-body docked into the maps and visually evaluated for agreement with the experimental density. All molecular visualizations, structural superpositions, and figure preparations were performed using UCSF ChimeraX [91,92]. Unless otherwise stated, default parameters were used in all prediction and analysis steps.

## DATA AVAILABILITY

The electron density maps and atomic coordinates have been deposited in the Electron Microscopy Data Bank (https://www.ebi.ac.uk/emdb/) and Protein Data Bank (https://www.rcsb.org) under accession codes EMD-75612, EMD-75635, EMD-75636, 11BX, 11DM, respectively. The complete ΦXacm4-11 genome sequence was submitted to the NCBI (BankIt) with the accession number PZ048659.

## ACKNOWLEDGMENTS AND FUNDING SOURCES

This work was funded by Fundação de Amparo à Pesquisa do Estado de São Paulo (FAPESP) grants #2017/17303-7 and #2021/10577-0 to C.S.F, and #2020/11189-0 and #2022/03018-7 to G.G.S. J.C.S. is the recipient of a CNPq Researcher Fellowship (#305274/2024-4) and E.E.L. acknowledges a FAPESP Postdoctoral Fellowship (#2019/12234-2). We thank Deyvid Amgarten for assistance with processing the bacteriophage sequencing data. We thank the Centro de Pesquisa em Biologia de Bactérias e Bacteriófagos (CEPID B_3_) for institutional support and for providing the collaborative and interdisciplinary environment that made this work possible. The authors thank LME/LNNano/CNPEM/MCTI for the access to the cryo-EM facility via proposals TEM-25042 and TEM-25942.

---

**Movie 1 Structural comparison of the nozzle regions of bacteriophages T7 and Xacm4-11.** The movie begins with independent rotations that highlight their overall architecture, followed by structural superposition to reveal a conserved core fold shared by the nozzle proteins from both phages despite the absence of detectable sequence homology. Additional domains unique to Xacm4-11 (ED1-ED4) are then highlighted to emphasize structural divergence. Final rocking motions reinforce similarities and differences between the two assemblies. This movie complements **Figure 5.**

**Movie 2 Sequential assembly of the Xacm4-11 portal-tail complex.** Visualization of the hierarchical assembly of the Xacm4-11 portal-tail complex. Structural components are incorporated sequentially to illustrate the organization and modular architecture of the portal (gp22), adaptor (gp29), nozzle (gp30 and gp31), and tail fibre protein gp36. The N-terminal cryo-EM-resolved and C-terminal AlphaFold3-predicted regions of the trimeric tail fibre protein gp36 are shown to highlight their integration within the complete structure. Surface rendering and sectional views reveal the internal channel of the assembled complex and emphasize its overall structural organization. This movie complements **Figure 7.**

