## Supplementary Figures for "Cryo-EM structure analysis of phage ΦXacm4-11 that infects the phytopathogen *Xanthomonas citri*"

**Figure S1**

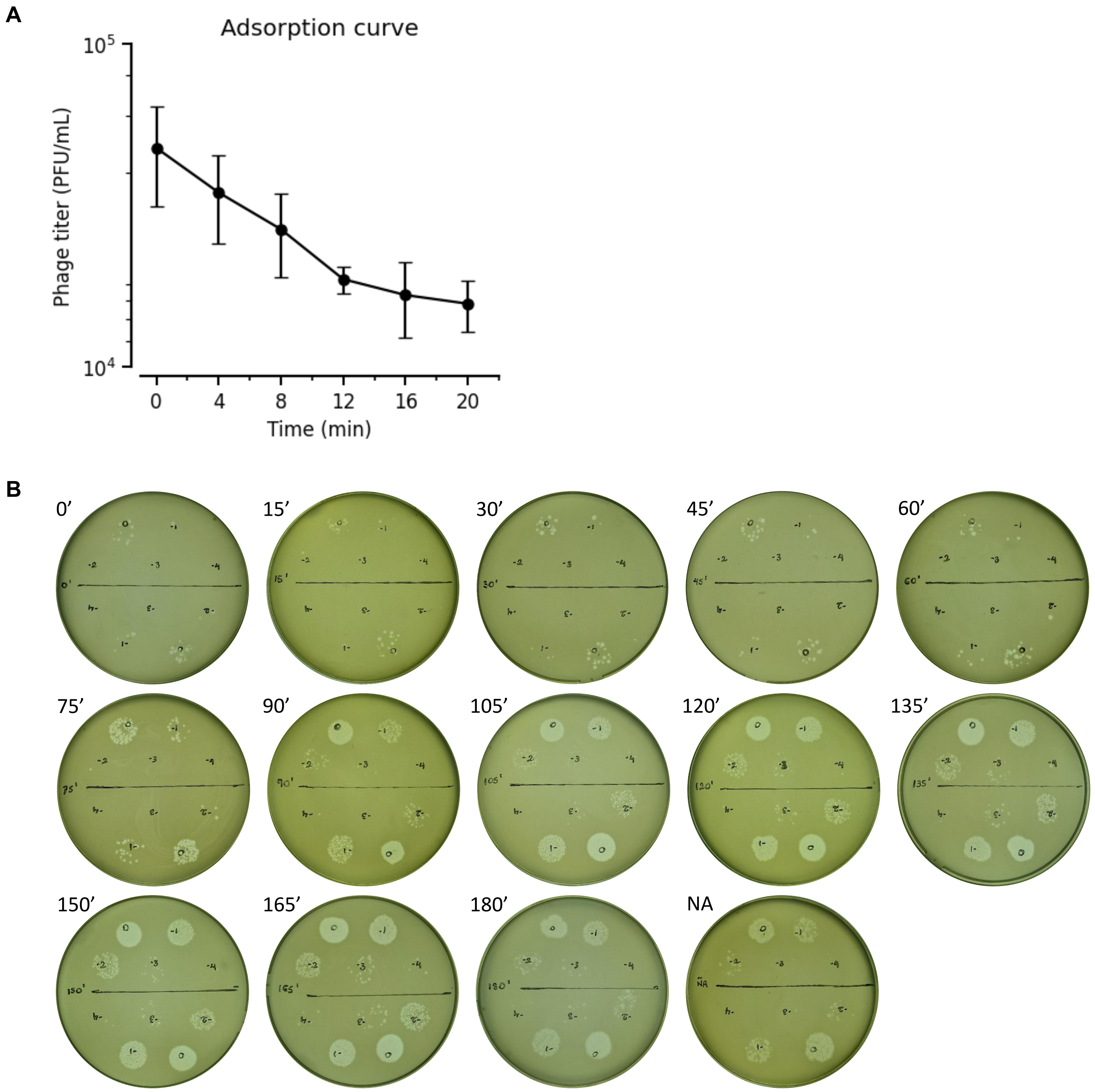


**Figure S1 | Representative plaque assays from kinetic experiments of ΦXacm4-11. (A)** Plaque assay-derived phage titres from the adsorption experiment, plotted as raw data underlying the curves shown in the main figure. **(B)** Plaque assay plates from the one-step growth experiment, with each plate corresponding to a specific time point. Titres were determined in duplicate on the same plate, and the experiment was performed in four independent biological replicates.

**Figure S2**

**A**

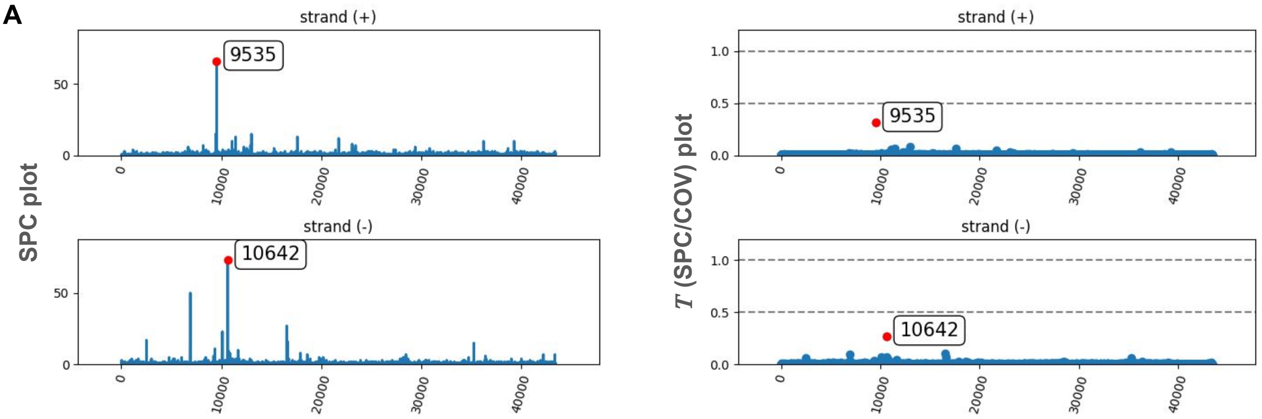


**Sequence coverage at termini position plot**

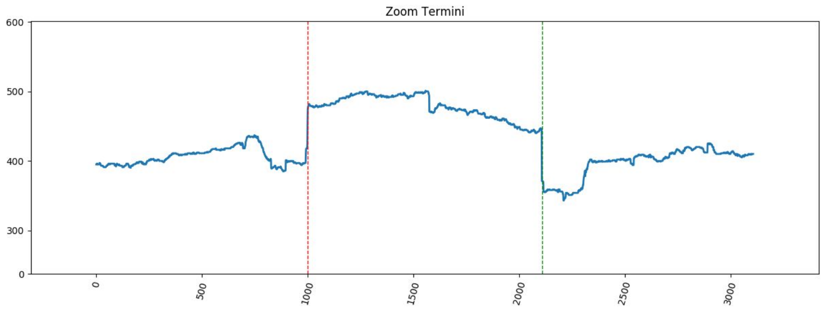


| Phage | | ΦXacm4-11 |
|---|---|---|
| Genome length (bp) | | 43,420 |
| Class | | DTR (long) |
| Termini position (bp) | Left (red) | 9,535 |
| | Right (green) | 10,642 |
| DTR size (bp) | | 1,108 |
| Ends | | Redundant |

**ΦXacm4-11 genome region containing the DTR predicted by the PhageTerm algorithm**

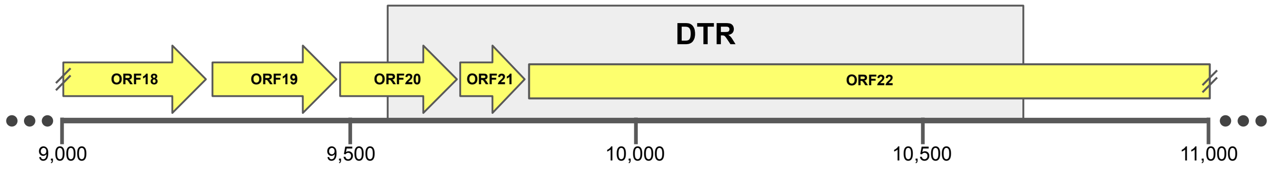


**Figure S2 | (A) Analysis of phage genome termini and DNA packaging mode using the PhageTerm algorithm [39].** The panel displays the outputs generated in the following order: (i) the starting position coverage (SPC) plot, showing the distribution of read start positions along the genome; (ii) the $T$ (SPC/COV) plot, highlighting the ratio between starting position coverage and overall sequence coverage; (iii) the sequence coverage profile at the predicted termini positions; (iv) a summary table reporting the phage name, genome length (bp), predicted class of DNA packaging, termini positions in base pairs (left terminus in red and right terminus in green), direct terminal repeat (DTR) size (bp), and end type; and (v) a schematic representation of the ΦXacm4-11 genomic region containing the predicted DTR by the algorithm, which starts near the 5′ end of ORF20 and extends through ORF22.

**Figure S2 (cont.)**

**B**

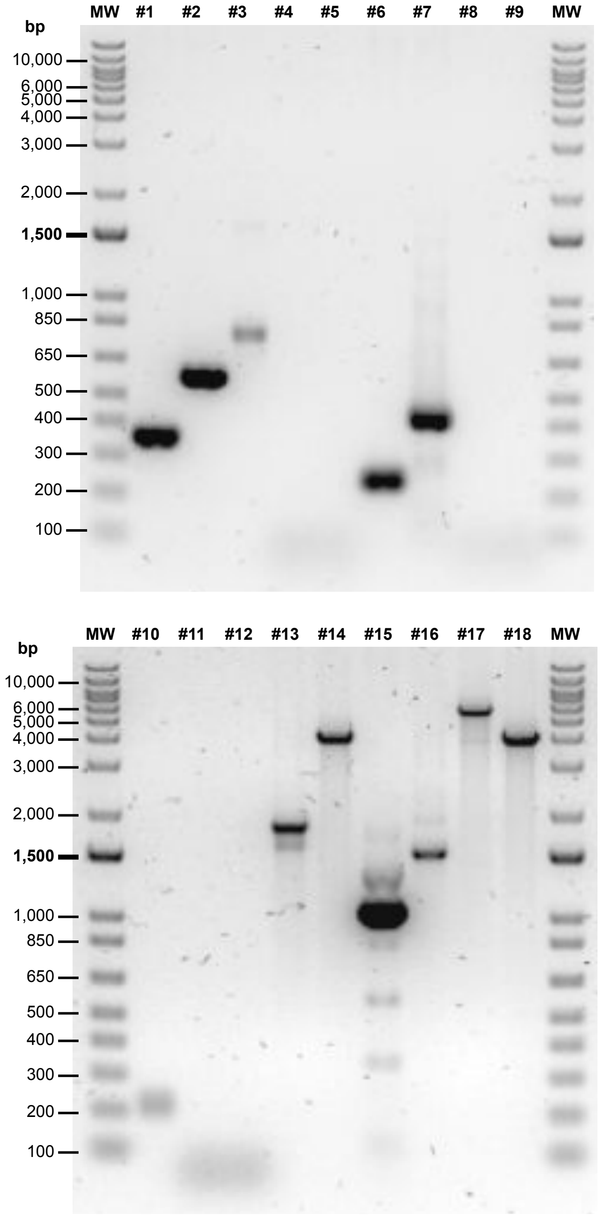


| Pair # | Primer Forward | Primer Reverse | Expected amplicon size (bp) if: | |
|---|---|---|---|---|
| | | | Circular | Linear (DTR) |
| **1** | ORF18_F | ORF18_R | 358 | 358 |
| **2** | ORF18_F | ORF19_R | 576 | 576 |
| **3** | ORF18_F | ORF20_R | 782 | 782 |
| **4** | ORF18_F | ORF22_R | 2,706 | No amplification |
| **5** | ORF18_F | ORF25_R | 4,823 | No amplification |
| **6** | ORF19_F | ORF19_R | 235 | 235 |
| **7** | ORF19_F | ORF20_R | 441 | 441 |
| **8** | ORF19_F | ORF22_R | 2,365 | No amplification |
| **9** | ORF19_F | ORF25_R | 4,482 | No amplification |
| **10** | ORF20_F | ORF20_R | 220 | 220 |
| **11** | ORF20_F | ORF22_R | 2,144 | No amplification |
| **12** | ORF20_F | ORF25_R | 4,261 | No amplification |
| **13** | ORF22_F | ORF22_R | 1,819 | 1,819 |
| **14** | ORF22_F | ORF25_R | 3,936 | 3,936 |
| **15** | ORF25_F | ORF25_R | 1,033 | 1,033 |
| **16** | ORF25_F | ORF26_R | 1,514 | 1,514 |
| **17** | ORF26_F | ORF31_R | 5,399 | 5,399 |
| **18** | ORF36_F | ORF39_R | 3,825 | 3,825 |

**Figure S2 (cont.) | (B) Experimental validation of phage genome linearity by PCR.** Agarose gel electrophoresis showing PCR amplification products obtained using primer pairs designed to probe the physical organization of the ΦXacm4-11 genome. The accompanying table summarizes the primers used and the expected amplicon sizes assuming either (i) a circular genome (without DTR) or (ii) a linear genome containing direct terminal repeats (with DTR) at the positions predicted by PhageTerm [39].

### Figure S3

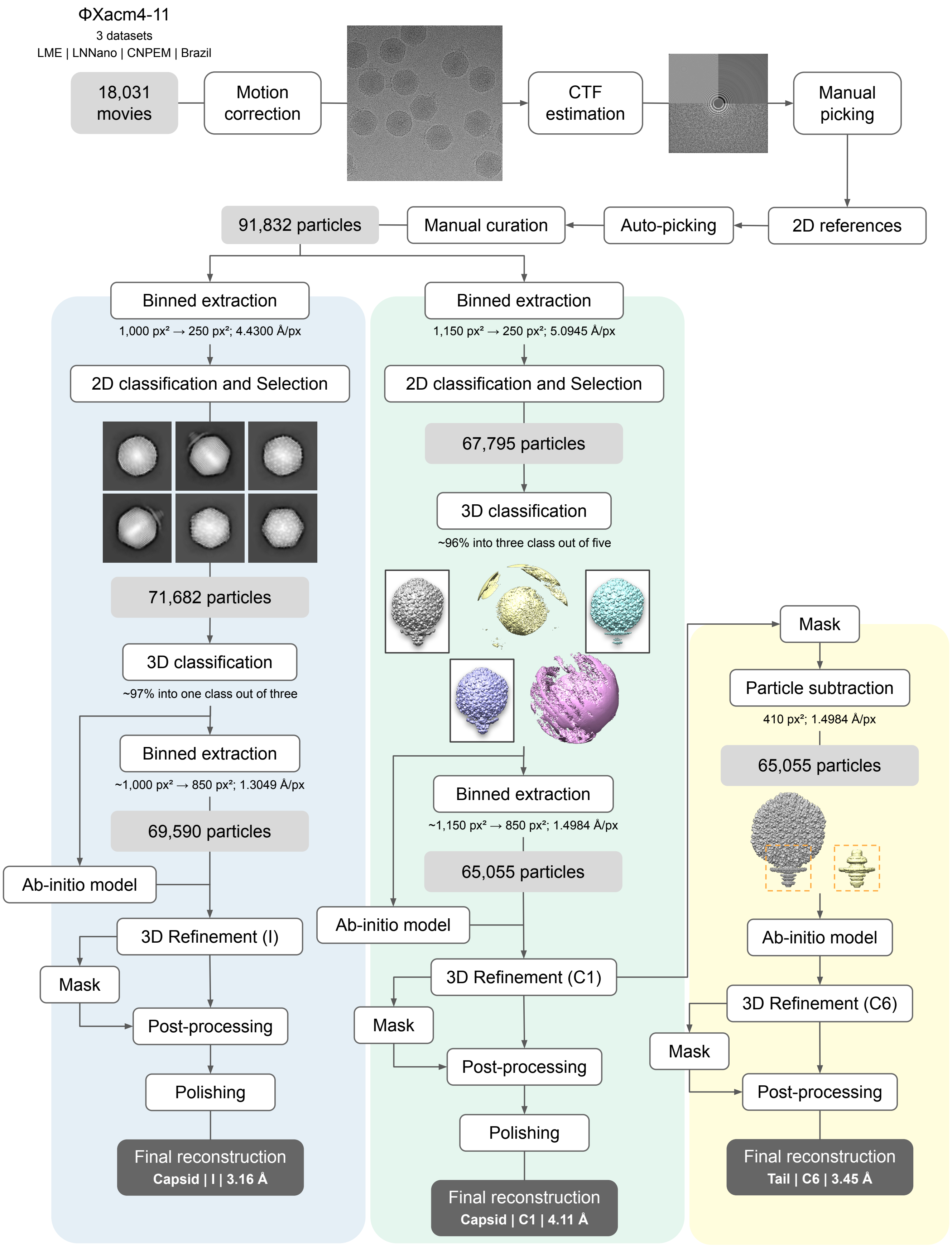


**Figure S3 | Cryo-electron microscopy data processing workflow.** Schematic overview of the cryo-EM data processing strategy implemented in RELION [42,80]. Three independent datasets were processed together following two parallel branches: one involving icosahedral symmetry (I) to obtain the symmetric capsid reconstruction, and a second branch based on C1 symmetry for asymmetric reconstruction. The latter pathway further diverged into a focused processing workflow, including particle subtraction and dedicated refinement steps, to resolve the structure of the tail region.

**Figure S4**

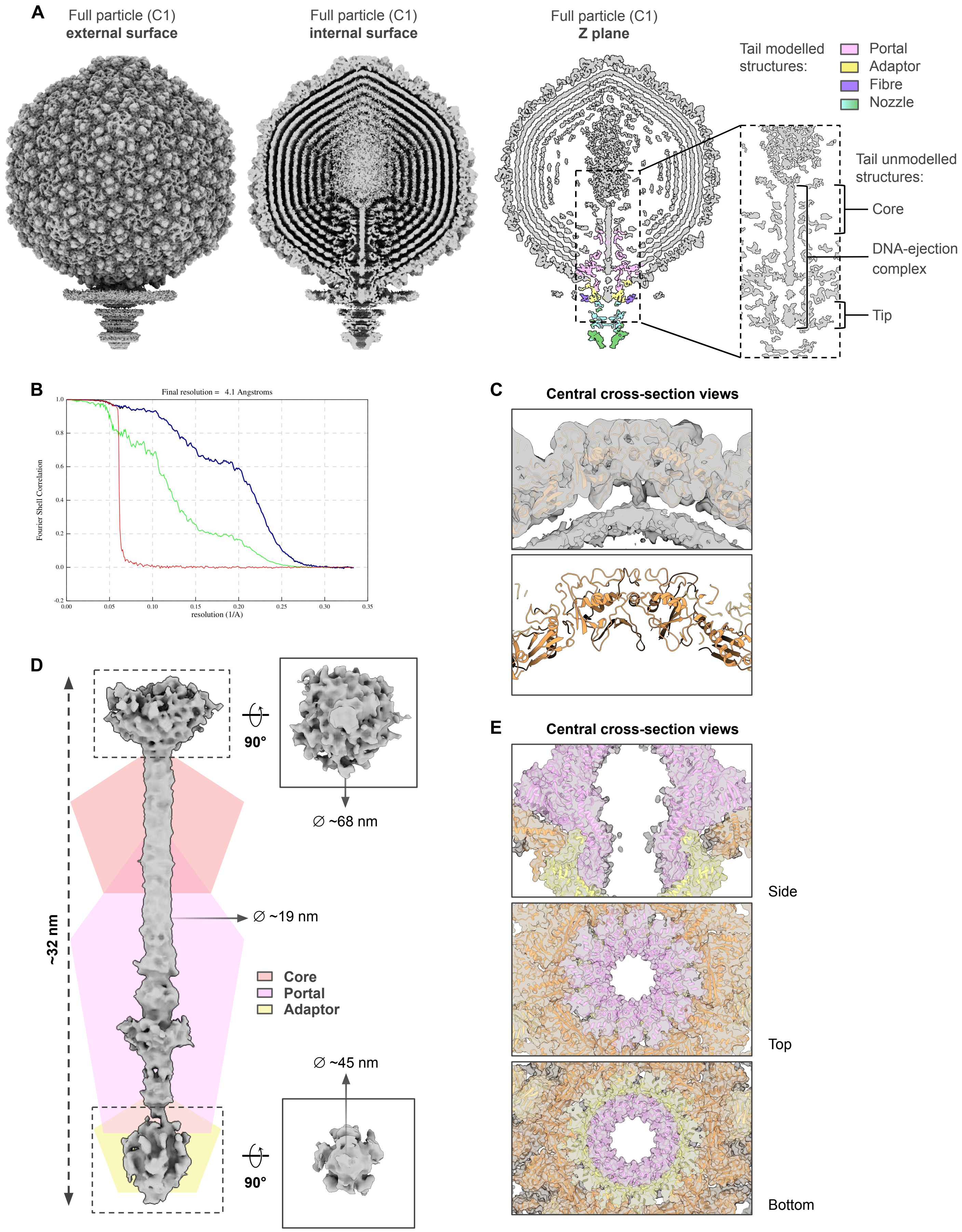


**Figure S4 | Structural features of the asymmetric reconstruction of ΦXacm4-11. (A)** Three views of the C1 cryo-EM reconstruction of ΦXacm4-11. *Left to right*: External surface rendering of the virion, a central slice revealing the internal organization of the particle and a slice view highlighting the tail region with modelled and unmodeled components indicated. **(B)** Gold-standard Fourier shell correlation (GSFSC) curve showing the resolution achieved for the asymmetric (C1) reconstruction of the phage virion. **(C)** Central slices at the interface between the capsid (MCP and CP) and the packaged double-stranded DNA. Two views are shown: one displaying the fitted protein models together with the corresponding cryo-EM density, and a second showing the protein models alone. **(D)** Cryo-EM map of the DNA-protein injection complex, with overall dimensions indicated along the length of the structure. Shaded regions denote the approximate locations of the core, portal, and adaptor proteins within the assembly. **(E)** Central slices through the interface between the tail (portal and adaptor proteins) and the special vertex of the capsid (MCP). Protein models are shown in ribbon and surface and visualized from side, top, and bottom perspectives. Animations of these regions are provided in **Movie S1**.

### Figure S5

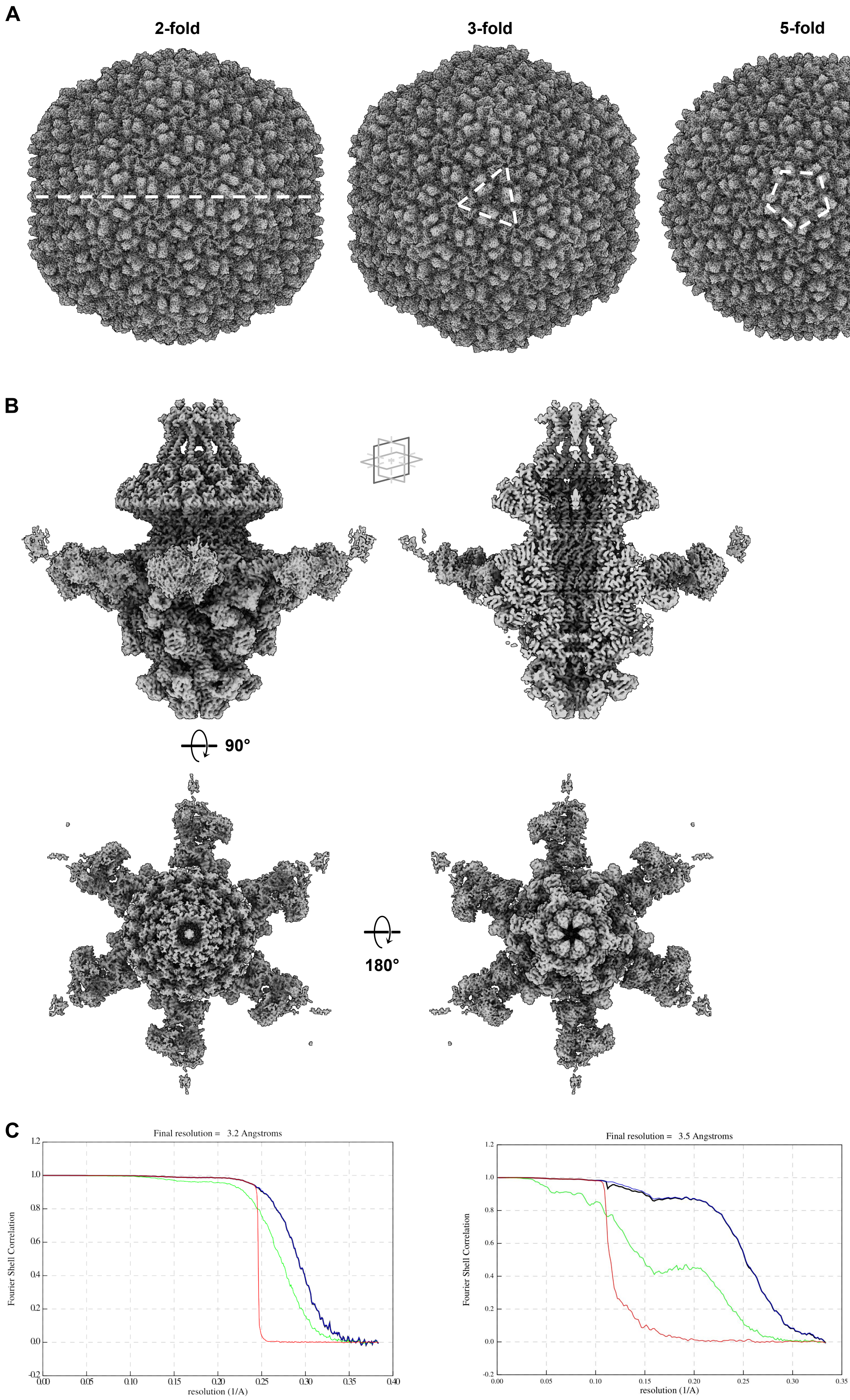


**Figure S5 | Symmetry-imposed reconstructions of the ΦXacm4-11 capsid and tail. (A)** Cryo-EM reconstruction map of the capsid refined under icosahedral (I) symmetry, displayed along the 2-, 3-, and 5-fold symmetry axes. **(B)** Cryo-EM reconstruction map of the tail refined with C6 symmetry. *Upper left to lower right*: Side full, side centrally-clipped, bottom and top views. **(C)** Gold-standard Fourier shell correlation (GSFSC) curves for the capsid and tail reconstructions, indicating overall resolutions of 3.2 Å and 3.5 Å, respectively.

**Figure S6**

**A**

```
T7 gp14            MCWAAAIPIAISGAQAISGQNAQAKMIAAQTAAGRRQAMEIMRQTNIQNADLSLQARSKL
Xacm4-11 gp33      MCEPITISTGAAWALGAAAAATAAATYVSVDAN--KKAGEANQQVAENNARLAADDAAAA
                   ** . :*. . : * . :.  : *   .:  *   ::* *  :*.  :** *: :  :

T7 gp14            EEAS-AELTSQNMQKVQAIGSIRAAIGESMLEGSSMDRIKRVTEGQFIREANMVTENYRR
Xacm4-11 gp33      QAMGDRESQAQTWRTRAIMGQQRAAIAANGIDAG------IGTPAEILGETALFGEVDQQ
                   :  .  *  :*. :.   :*. ****. . ::..        * .::: *: :. *  ::

T7 gp14            DYQAIFAQQLGGTQSAASQIDEIYKSEQKQKSKLQMVLDPLAIMGSSAASAYASGAFDSK
Xacm4-11 gp33      AIRLNTARTAWG------------FNSQVRNIQNQAGIDRFNTKAKGTATVLSG--ISSI
                     :   *:   *             ..* :: : *  :* :   ...:*:. :.  :.*

T7 gp14            STTKAPIVAAKGTKTGR
Xacm4-11 gp33      ASSGAGFYG--------
                   ::: * : .
```

**Figure S6 | (A)** ClustalW sequence alignment between the bacteriophage T7 core protein gp14 and its predicted homolog in ΦXacm4-11, gp33.

**Figure S6 (cont.)**

**B**

```
T7 gp15            -------MSKIESALQAAQPGLSRLRGGAGGMGYRAATTQAEQPRSSLLDTIGRFAKAGA
Xacm4-11 gp34      MATIIPRVTGPQVEVQQGPQVRNTARVDDSGAQALAGAIGSFARPALEYAQREQERNDTT
                          ::  :  :* .    .  * . .*    *.:  :    :       :  :  :

T7 gp15            DMYTAKEQRARDLADERSNEIIRKLTPEQRREALNNGTLLYQDDPYAMEALRVKTGRNAA
Xacm4-11 gp34      AVMAARKQLSDWEASTFNPENPEGINKYKGKNALGADEVLLPDLDKTVGSISASLTPRQR
                    : :*::* :   *.  . *  . :.  : ::**. . :*  *   :: :: ..   .

T7 gp15            YLVDDDVMQKIKEGVFRTREEMEEYRHSRLQEGAKVYAEQFGIDPEDVDYQRGFNGDITE
Xacm4-11 gp34      QQFEGVAMNFRDSVAGRLNGYMDREHSGYISAEQKAAVDNLGADAVSAGVAGDFSRQDRV
                     .:. .*:  .. . * .  *:. : . :.   *. .:::* *. ...   .*. :

T7 gp15            RNISLYGAHDNFLSQQAQKGAIMNSRVELNGVLQDPDMLRRPDSADFFEKYIDNGLVTGA
Xacm4-11 gp34      ANELLAMNRARRQADGLGEELIKAEERGLVSSVRSQTIEGMATARPFEAQAYFERYADQM
                    *  *   : .  ::   :  *  ..  * . ::.  :   . :  *  :   :  .

T7 gp15            IPSDAQATQLISQAFSDASSRAGGADFLMRVGDKKVTLNGATTTYRELIGEEQWNALMVT
Xacm4-11 gp34      TPEDRAKVDRLLYPVVSDAAAQDDADAILAGAQPAAYRDPGQRGRGPSPAVAKILDEEAD
                    *.*   .: :  .. . ::  ..** ::  .:  .  : .        .  :     .

T7 gp15            AQRSQFETDAKLNEQYRLKINSALNQEDPRTAWEMLQGIKAELDKVQPDEQMTPQREWLI
Xacm4-11 gp34      AAGVPREYLYALAEQESGFNPGAVNPEVLDDGDNATGLFQYRATSAKGFDRTDARASARA
                   *     *    * **      .*:* *    . :    :: .  ..:  ::  .: .

T7 gp15            SAQEQVQNQMNAWTKAQAKALDDSMKSMNKLDVIDKQFQKRINGEWVSTDFKDMPVNENT
Xacm4-11 gp34      AAREFAQRAKTGGVEFAVAAHFAGDGGAEAVVQRGRRAENPKTARYVEEVMGRASRWRGD
                   :*:* .*.  .. .:  . *   .  . : :   .:: ::  ...:*.  :   .  ..

T7 gp15            GEFKHSDMVNYANKKLAEIDSMDIPDGAKDAMKLKYLQADSKDGAFRTAIGTMVTDAGQE
Xacm4-11 gp34      APATQVGRTAPPSEADAIARAQAIPDPRRRAAVVSKIRERFQLSDLRRQE---EDRAASE
                   .  .: . .  ..:  *   :  ***  : *  :. ::   : . :*         *..*

T7 gp15            WSAAVINGKLPERTPAMDALRRIRNADPQLIAALYPDQAELFLTMDMMDKQGIDPQVILD
Xacm4-11 gp34      GAYTAINQAANPNAPLREILG--------AQVYAWAERKGHLNTLENLRKNKVAGTFTQD
                    : :.**     .:*  : *           .  :.::   : *:: : *: :   .  *

T7 gp15            ADRLTVKRSKEQRFEDDKAFESALNASKAPEIARMPASLRESARKIYDSVKYRSGNESMA
Xacm4-11 gp34      DAVLVESLEREAVLSPQTFARRDIHALADRISTQTLGELIGKQKQANDPSKR--ADWANQ
                      *. . .:*  :. :.  .  ::*      ::  ..*  . ::  *. *   .: :

T7 gp15            MEQMTKFLKESTYTFTGDDVDGDTVGVIPKNMMQVNSDPKSWEQGRDILEEARKGIIASN
Xacm4-11 gp34      SQRIESGLRILGLDEAGDNRNAEGEFVAKPPKKKLDERKVQRAQFAQVYREAERAFIQR-
                    ::: . *:      :**: :.:   *      :::.   .  *  :: .**.:.:*

T7 gp15            PWITNKQLTMYSQGDSIYLMDTTGQVRVRYDKELLSKVWSENQKKLEEKAREKALADVNK
Xacm4-11 gp34      ----HGKDPTPAEADSILRAVVRNVATDIPGKLSRAGAIEGFGAALPAADRSEIISDFRA
                       : : .  ::.***    . . .    .*   : . .     *    *.: ::*..

T7 gp15            RAPIVAATKAREAAAKRVREKRKQTPKFIYGRKE
Xacm4-11 gp34      STGREPTEAEIVRIGARYRMQSNSAGSN------
                    :   .:       . * * : :.: .
```

**Figure S6 | (B)** ClustalW sequence alignment between the bacteriophage T7 core protein gp15 and its predicted homolog in ΦXacm4-11, gp34.

**Figure S6 (cont.)**

**C**

```
T7 gp16            MDKYDKNVPSDYDGLFQKAADANGVSYDLLRKVAWTESRFVPTAKSKTG----PLGMMQF
Xacm4-11 gp35      MGKYSEIIAATQAEDDQARARAAGITAPQVEPDAMGRAMRLADQRGLPPGVVADNLPEYD
                   *.**.: :.:      *  * * *::   :.  *  .:  :.  :. .

T7 gp16            TKATAKALGLRVTDGPDDDRLNPELAINAAAKQLAGLVG---------------------
Xacm4-11 gp35      QQQQLDALGDAASASPVLARWLGDPNHTALSKDDTGVLGGLGRQFNSGLLSAKETLPWLP
                    :   .***  .: .*   *   :   .* :*: :*::*

T7 gp16            ----KFDGD-ELKAALAYNQGEGRLGNPQLEAYSKGDFASISEEGRN-------------
Xacm4-11 gp35      LPQSALDLDKERKAAAARGPQMMASPNPSWFSELVGKFQSGWEQGKAGTGLLPMPNLGSA
                        :* * * *** * .       **.  :   *.* *  *:*:

T7 gp16            ------------------------------------------------------------
Xacm4-11 gp35      DSRSVEEQVARGLGYQSAADYSTAQTAVYVADQQRRAQAADVETAATAAGFKEIADAEKT

T7 gp16            -----YMRNLLDVAKSPMAGQLETFGGITPKGKG---IPAEVGLAGIGHKQKVTQELPES
Xacm4-11 gp35      KDTGKIARAVVTNPRAVMAVVAQSLGNTAPQLALSAAMPASRIVAGVTAGTGSFATEQGM
                          * ::  .:: **   :::*. :*:      :**.  :**:

T7 gp16            TSFDVKGIEQEATAKPFAKDFWETHGETLDEYNSRSTFFGFKNAAEAELSNSVAG-----
Xacm4-11 gp35      TILDVMGDAGVETSDPAAVSAFLRNKPAMEKARQKALTRGVAVGTFDALTAGFAGRLIAN
                   * :** *     *:.* * . :  :  :::: ..::   *.  .:   *: ..**

T7 gp16            ------------------------------------------------------------
Xacm4-11 gp35      SRRTAVSIGGRVAGEAAVQAGGEIQDAITAGLAEAQQQESRENGPAGQVYDDVVGQLLSR

T7 gp16            ------------------------------------------------------------
Xacm4-11 gp35      QDEQLARQNAAVVQSVYRNLAERVGTDAWTLYQQFKLRIPGATTDTRARPRGVDIDVDPF

T7 gp16            -MAFRAGRLDNGFDVFKDTITPTRWNS---------------------------------
Xacm4-11 gp35      LDALRSGKMPTDREIYGDTLVSALHAAGGLRDSGGELANMDAAKARPGLVNNLAGMSLDD
                     *:*:*:: .. ::: **:..:   :

T7 gp16            --------------------------------------------HIWTPEELEKIRTEVK
Xacm4-11 gp35      ALVWAYQEGFITQAPTNQQEGSYDADAPDINTLLDLLAQDLGGNPVYRPAAMNAERAAFR
                                                                :: *  ::  *: .:

T7 gp16            N-----------------------------------------------------------
Xacm4-11 gp35      DRAQSLQDELDQRGIDLQQADNATARQALGYANRTLEQAPQPVETDGVDYAVRDLVTGAS
                   :

T7 gp16            ------------------------------------------------PAYINVVTGGSP
Xacm4-11 gp35      GRPESAATRALVDQAVADGDLAKMLRGIAAQEGIDADQAALALRLAEITPDLNVTMVPAP
                                                                   .. :**.   :*

T7 gp16            ENLDDLIKLANEN-----------------------------------------------
Xacm4-11 gp35      ANANAAGIYNNTTNEIWIHQAVPSVVLHEVLHGVTSAMMTSPTLRRANPTVARAVAEFND
                    * :      * .

T7 gp16            ---FENDSRAAEAGLGAKLSAGIIGAGVDP------------------------------
Xacm4-11 gp35      LLGAAQAHFAGMETDGADVPAALRATLQDPRGPLTNIKELLTYGMTDKRFQEWLATVPAP
                        :   *.    **.:.*.:  .:  **

T7 gp16            --------------LSYVPMVGVTGKG---------------------------------
Xacm4-11 gp35      AGREEARTAWQWFKDAIATMLGVTGKERTALDALIESTSDLVDFAQANPRAANFAQMSEA
                                  : ..*:*****
```

**Figure S6 (cont.)**

**C (cont.)**

```
T7 gp16            --------------------FKLINKALVVGAESAALNVASEGLRTSVAGGDADYAGAAL
Xacm4-11 gp35      NRLGRPVADAAAMEGGPAEAFRGVTREQFLGEPRITANSNAAELRPRALRELDDVEAVPF
                                       *: :.:  .:*    : *  :  **. .     *  ...:

T7 gp16            GGFVFGAGMSAISDAVAAGLKRSKPEAEFDNEFIGPMMR---------------------
Xacm4-11 gp35      GDGDLTAKYSDGSAAVFDGDKVVASYNFGDTLVVDKAYRRRGIGEELVYQWRTRNPQASV
                   *.  : *  *  * **  * *    .    *. .:.   *

T7 gp16            ------------------------------------------LEARETARNANSADLSRM
Xacm4-11 gp35      ARERTKASQALQEKVWDRIQSELAADPRTLNQGGENPRGAVTFEGSPGARVFNIELLKGM
                                                             :*.   **  *   *. *

T7 gp16            NTENMKFEGEHNGVPYEDLPTERG---------AVVLHDGSVLSASNPINPKTLKEFSEV
Xacm4-11 gp35      DASTFMHEMGHVYLEVLNDLAARDGAPAQIVNDMATLNAWLGRDAGAAFTVDQHEQFARG
                   ::.: .*  *  :   :  : *.          ..*:     .*. .:. .  ::*:.

T7 gp16            DPEKAARGIKLAGFTEIGLKTLGSDDADIRRVAIDLVRSPTGMQSGASGKFGATASDIHE
Xacm4-11 gp35      FEAYLREGRAPSSALRRTFAAFKVWLTALYRSVRSLNVELTDDVRNVMDRIVASDAEIED
                         .*   :.  .  : ::    : : * . .*  . *.   .. .:: *: ::*.:

T7 gp16            RLHGTDQRTYNDLYKAMSDAMKDPEFSTGGAKMSREETRYTIYRRAALAIERP-------
Xacm4-11 gp35      ARAGQYQGALIADGMAVGMTFEQLQAYNEAVNAARADAEATVAAEVLLAERREQQAWYNR
                      *  * :      *:. :::: :  . ..: :* ::. *:  .. ** .*

T7 gp16            ------------------------------------------------------------
Xacm4-11 gp35      EKREVRAKVLEEVRNLPVYRAQRLLRNGKLPNGDLAPDELRVKLSKDELLDLYGQSFLRN


T7 gp16            --------------------------ELQKALTPSERIVMDIIKRHFDTKRELMENPAIF
Xacm4-11 gp35      LVGMYSVEGGVRADEAAAIYGFGSGRELVESLVNAPRLADAVASETDSRMKDRYPDPMTD
                                             ** ::*. : *:.  : ..  .  ::   :*

T7 gp16            GNTKAVSIFPESR---------HKGTYVPHVYDRHAKALMIQR-----------------
Xacm4-11 gp35      GTLPDRAMIAAHRDRQADVMVREIRALEQHVSGRQVSQAAVIKGVARQIIQDKKLRHLQP
                   *.    :::.  *          .  :   ** .*:..   : :

T7 gp16            ---------YGAEGLQEGIARSWMNSYVSR-----------PEVKARVDEMLKELHGVKE
Xacm4-11 gp35      ATYRSAEARAGREAFEAAAKQDWGSALAARRRQLLNFELFREAVRARDEAARTAKYLAKF
                             * *.:: .  :.* .: .:*             *:** :   .  : .*

T7 gp16            VTPEMVEKYAMDKAYGISHSD----QFTNSSIIEENIEGLVGIENNSFLEARNLFDS---
Xacm4-11 gp35      SETKTRARLGKAGADYLDQVDGLLDRFDFRKISDKAADRRSSLATWIEVQAQKGIDVNLP
                     .:   : .   *  :.: *    :*   .* ::  :   .: .   ::*:: :*

T7 gp16            ----DLSITMPDGQQFSVNDLRDFDMFRIMPAYDRRVNGDIAIMGSTGKTTKELKDEILA
Xacm4-11 gp35      PKIMDEAFTIPYREMTVADLATLRDAIRSIDHLAR-LKGKLLLAGEVRDAAEIDAMAAS
                       * ::*:*  :   .:     * :* :    * ::*.: : *.. .:::       :

T7 gp16            LKAKAEGDGKKTG---------------EVHALMDTVKILTGRARRNQDTVWETSLRAIN
Xacm4-11 gp35      LAAAHAARPVSTGDRGPKDKLRQAFMQGRVIQATATDIAREL DGFKDQGAIWMNTVGVMR
                   * *   .   .**                 .*     *        . ::*.::* .:: .:.

T7 gp16            DLG-----------------------------------------FFAKNAYMGAQNIT
Xacm4-11 gp35      DAVNNRLNPALQQAQDELAQVYVKHYSKAEIRGFSERVPMPEVNGDLWSKSRLLGLALNW
                   *                                              :::*.  :*

T7 gp16            EIAGMIVTGNVRALGHGIPILRDTLYKSK-------------------------------
Xacm4-11 gp35      GNAGNREAILTQARGRMSPEQVNALLSKLDSRDWAFVEDVWRLIDAQWPAIAEAQKRRTG
                     **   :  .:* *:  *   ::* ..
```

**Figure S6 (cont.)**

**C (cont.)**

```
T7 gp16            -------------PVSAKELKELHASLFGKEVDQLIRPKRADIVQRLREATDTGPAVANI
Xacm4-11 gp35      LVPERVQASAFTVQTSDGKTLQIPGGYYPLKYESDSVKTMKDEADDFYNSIRTGRTAKAA
                                 .*  :  :: .. :  : :.    .  * .: : ::  ** :.

T7 gp16            VGTLKYSTQELAARSPWTKLLNGTTNYLLDAARQGMLGDVISATLTGKTTRWEKEGFLRG
Xacm4-11 gp35      TRNGHTIERVGSGGRTVRLDTGVVQQHLRDVLRDVHLGDAVNYVHNVLNGQEFKEAVDST
                   . . :   :  :.  .     . . ::* *. *:  ***.:. . .  . :  **..

T7 gp16            ASVTPEQMAGIKSLIKEHMVRGEDGKFTVKDKQAFSMDPRAMDLWRLADKVADEAMLRPH
Xacm4-11 gp35      GTQEYRQALEVWLKDAAAGEIGPRVWHERAMRAARQNFTASVLTFKVTSALLQLSGVVPT
                   .:   .*   :          *    .    : * .  . ::  ::::. : : : : *

T7 gp16            KVSLQDSHAFGALGKMVMQFKSFTIKS------------LNSKFLRTFYDGYKNNRAIDA
Xacm4-11 gp35      IVTIGQGHTMAGIGQYLGKPRAMTRYVREASPYMDSRLRTHIEAVQTVMDAEAGRFAAGK
                    *:: :.*::..:*: : :::*                : : ::*. *.  .. * .

T7 gp16            ALSIITSMGLAGGFYAMAAHVKAYALPKEKRKEYLERALDPTMIAHAALSRSSQLGAPLA
Xacm4-11 gp35      AASIRFGYWMIGRVQGLVDTVTWLAAEQAGMAKFDNDVARARAYADDVVTRAQGSGEFID
                   * **  .  : * . .:.  *.  *  :    :: : .  .   *. .::*:.  *  :

T7 gp16            MVDLVGGVLG--------------FESSKMARSTILPKDTVKERDPNKPYTSREVMGAMG
Xacm4-11 gp35      KSPLQRGTLGDNVRQTEWIKATTALQGYMIAKGNLAYEQTRKANLRNPRQAMKWAADMVM
                      *  *.**              ::.  :*:..:  ::* * .  *   : : . . :

T7 gp16            SNLLEQMPSAGFVANVGATLMNAAGVVNSPNKATEQDFMTGLMNSTKELVPNDPLTQQLV
Xacm4-11 gp35      LFSVEGLLTAALTAKLPKDDDDDGLWDDLGEWAIKDALSTFFGVIPGGGVLVDQFRGYDS
                      :* : :*.:.*::     : .   :  : * :: : * :   .   *  * :

T7 gp16            LKIYEANGVNLRERRK--------------------------------------------
Xacm4-11 gp35      SGVVAGAWRAYAELLEKVTPGEDGEVNLDKGVAKAAVSAAGVTLGLPSTQVNKTIDAIAA
                     :  .      *  :

T7 gp16            ---------------------
Xacm4-11 gp35      RADGRDVSPYEYLTGPKKESK
```

**Figure S6 | (C)** ClustalW sequence alignment between the bacteriophage T7 core protein gp16 and its predicted homolog in ΦXacm4-11, gp35.

### Figure S7

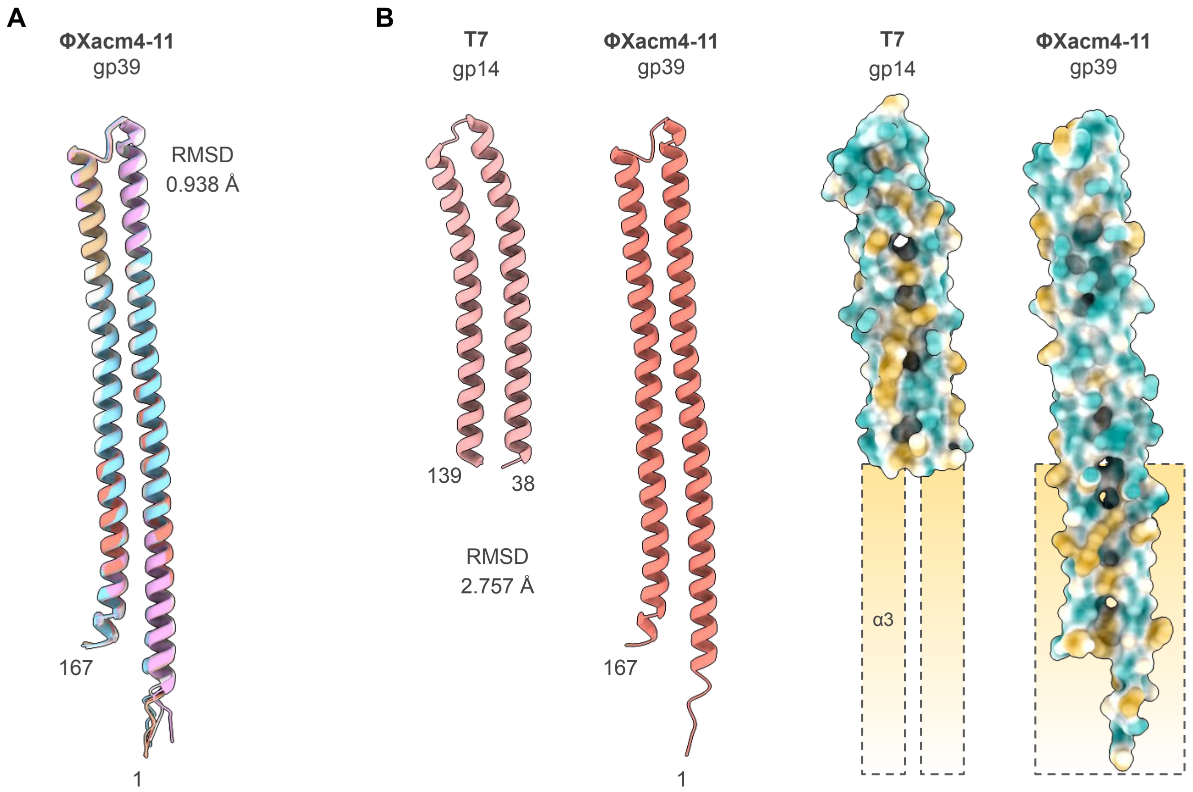


**Figure S7 | Structural comparison of the ΦXacm4-11 core protein gp33 and its T7 homolog. (A)** Superposition of the five AlphaFold3-predicted models of gp33, showing a high degree of structural consistency. **(B)** Structural comparison between one representative AlphaFold3 model of ΦXacm4-11 gp33 and the cryo-EM-derived structure of the bacteriophage T7 core protein gp14 (PDB: 7ey7). *Left to right*: Ribbon representations and surface representations colored by hydrophobicity. In the T7 gp14 structure, the membrane-interacting hydrophobic region previously reported by the authors [73], involving the N-terminus plus the C-terminal region formed by helix α3, is highlighted. A comparable hydrophobic region is observed in ΦXacm4-11 gp33, as indicated by the hydrophobic surface coloring. RMSD values among all atoms are indicated in each case.

**Figure S8**

**A**

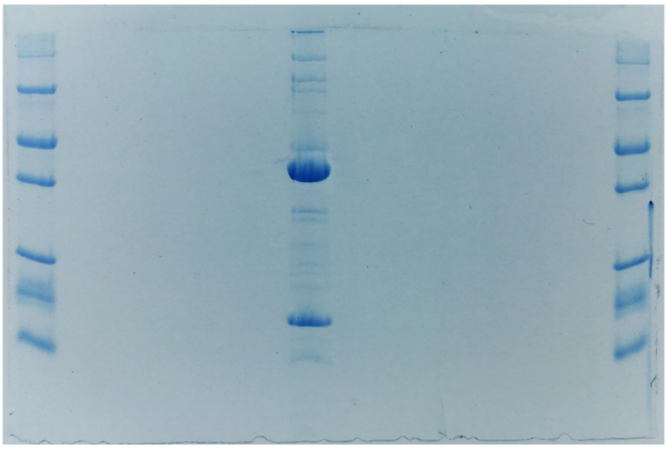

**B**

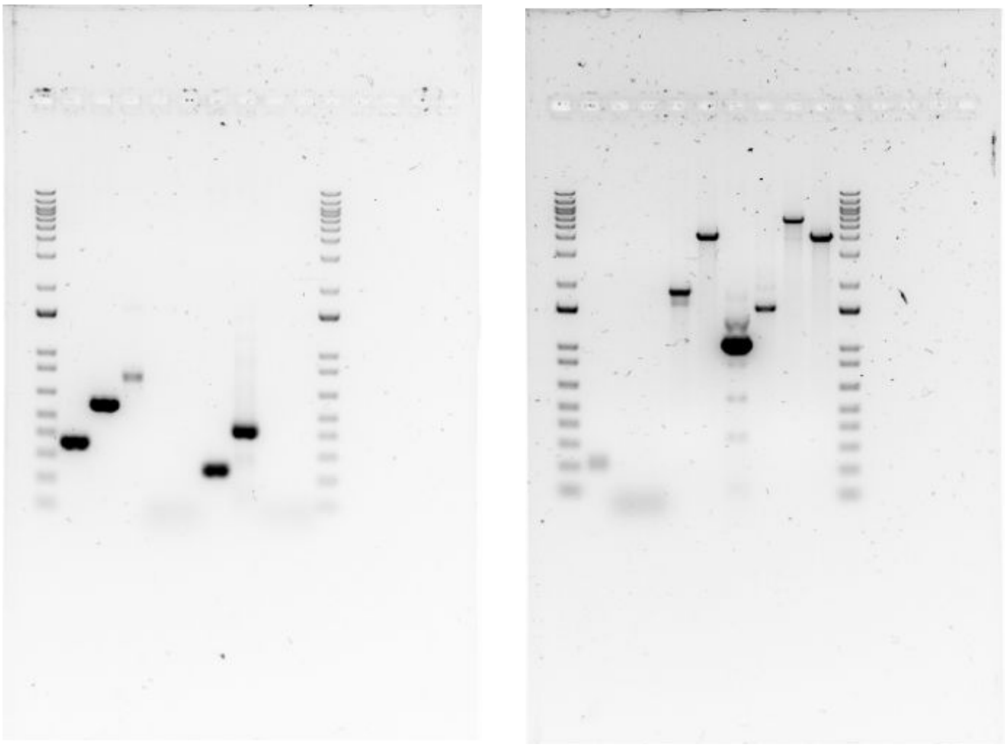

**Figure S8 | Source gels. (A)** Full SDS-PAGE gel used to generate the cropped panel shown in Figure 1D. **(B)** Uncropped 1% agarose gels corresponding to the PCR assays used to assemble Figure S2B.
