## Supplementary Tables for "Cryo-EM structure analysis of phage ΦXacm4-11 that infects the phytopathogen *Xanthomonas citri*"

| **Table S1 \| List predicted gene class and function in ΦXacm4-11, and homologues in CP2.** | | | |
|---|---|---|---|
| **Gene class** | **Predicted gene function** | **Name in ΦXacm4-11** | **Homolog in CP2** |
| II | Hypothetical protein | gp1 | |
| | Hypothetical protein | gp2 | |
| | Hypothetical protein | gp3 | ORF2 |
| | Hypothetical protein | gp4 | |
| | HNH Endonuclease | gp5 | |
| | Hypothetical protein | gp6 | |
| | RecA-dependent nuclease | gp7 | |
| | RusA endodeoxynuclease | gp8 | |
| | Lysozyme_like superfamily | gp9 | |
| | Hypothetical protein | gp10 | ORF6 |
| | Hypothetical protein | gp11 | ORF7 |
| | HNH endonuclease | gp11b | |
| | Hypothetical protein | gp12 | ORF8 |
| | Hypothetical protein | gp13 | ORF9 |
| | Hypothetical protein | gp14 | |
| | Terminase small subunit | gp15 | ORF10 |
| | Hypothetical protein | gp16 | |
| | Terminase large subunit | gp17 | ORF11 |
| | Hypothetical protein | gp18 | ORF12 |
| | Hypothetical protein | gp19 | |
| | Hypothetical protein | gp20 | |
| | Hypothetical protein | gp21 | |
| III | **Portal protein** | gp22 | ORF13 |
| | Hypothetical protein | gp23 | |
| | Putative protease | gp24 | ORF14 |
| | **Major capsid protein** | gp25 | ORF15 |
| | **Cement protein** | gp26 | ORF16 |
| | Hypothetical protein | gp27 | ORF17 |
| | Endonuclease | gp28 | |
| | **Adaptor protein** | gp29 | ORF18 |
| | **Tail tubular protein** | gp30 | ORF19 |
| | **Nozzle protein** | gp31 | ORF20 |
| | Internal virion protein | gp32 | ORF21 |
| | Internal virion protein | gp33 | ORF22 |
| | Lysozyme-like protein / Endolysin | gp34 | ORF23 |
| | Hypothetical protein | gp35 | ORF24 |
| | **Tail fibre protein** | gp36 | ORF25 |
| | Tail fibre protein | gp37 | ORF26 |
| | Tail fibre protein | gp38 | ORF27 |
| | Tail fibre protein | gp39 | ORF28 |
| I | Hypothetical protein | gp40 | |
| | Hypothetical protein | gp41 | |
| | Hypothetical protein | gp42 | ORF30 |
| | Hypothetical protein | gp43 | ORF31 |
| | Anti-CRISPR Vcrx091 protein | gp44 | ORF32 |
| | Exonuclease | gp45 | ORF34 |
| | HNH endonuclease | gp45b | |
| | Hypothetical protein | gp46 | |
| | ssDNA-binding protein | gp47 | ORF35 |
| | Homing endonuclease | gp47b | |

| **Table S1 (cont.) \|** List predicted gene class and function in ΦXacm4-11, and homologues in CP2. | | | |
|---|---|---|---|
| **Gene class** | **Predicted gene function** | **Name in ΦXacm4-11** | **Homolog in CP2** |
| I | Hypothetical protein | gp48 | |
| | Hypothetical protein | gp49 | |
| | Hypothetical protein | gp50 | ORF36 |
| | Hypothetical protein | gp51 | ORF37 |
| | Hypothetical protein | gp52 | ORF38 |
| | Hypothetical protein | gp53 | |
| | Hypothetical protein | gp54 | |
| | Hypothetical protein | gp55 | |
| | Hypothetical protein | gp56 | |
| II | Hypothetical protein | gp57 | |
| | Hypothetical protein | gp58 | ORF39 |
| | Hypothetical protein | gp59 | |
| | Hypothetical protein | gp60 | ORF40 |

ORFs shown in bold correspond to proteins for which structures were modelled in the cryo-EM maps.

**Table S2 |** Statistics of data collection, 3D reconstructions, and model building and refinement.

| | Bacteriophage ΦXacm4-11 mature virions | |
|---|---|---|
| **Data collection** | | |
| **Facility** | LME, LNNano, CNPEM | |
| **Microscope** | FEI Titan Krios | |
| **Detector** | Falcon 3EC | |
| **Magnification** | ×59,000 | |
| **Voltage (kV)** | 300 | |
| **Electron exposure ($e^{-}$/Å$^2$)** | 30 | |
| **Pixel size at collection (Å)** | 1.1075 | |
| **Defocus range (μm)** | -0.5 to -2.0 (step 0.25) | |
| **Frames per movie** | 20 | |
| **Movies** | 18,031 | |
| | **ΦXacm4-11 capsid** | **ΦXacm4-11 tail** |
| **3D Reconstruction** | | |
| **PDB entry code** | XXXX | XXXX |
| **PDB entry code** | XXXX | XXXX |
| **Particles per reconstruction** | 69,590 | 65,055 |
| **Symmetry** | I | C6 |
| **Box size (pixels)** | 850 (1.3049 Å/px) | 410 (1.4984 Å/px) |
| ***B* factor (Å$^2$)** | -129.74 | -82.57 |
| **Map resolution (Å)** | 3.16 | 3.45 |
| **FSC threshold** | 0.143 | |
| **Initial model used** | *de novo* | *de novo* |
| **Model composition** | | |
| **Peptide chains** | 840 | 54 |
| **Protein residues** | 203,280 | 17,754 |
| **RMS deviations** | | |
| **Bond lengths (Å)** | 0.003 | 0.004 |
| **Bond angles (°)** | 0.558 | 0.516 |
| **Ramachandran plot** | | |
| **Favored (%)** | 94.68 | 93.53 |
| **Allowed (%)** | 5.32 | 6.47 |
| **Outliers (%)** | 0.00 | 0.00 |
| **Validation** | | |
| **MolProbity score** | 1.50 | 1.59 |
| **Clashscore** | 3.47 | 3.76 |
| **CaBLAM outliers (%)** | 3.12 | 4.11 |
| **Rotamer outliers (%)** | 0.43 | 0.11 |

**Table S3 |** Positions of the MCP and CP subunits at the hexon-hexon and hexon-penton interfaces.

| Protein | Symmetry | Chain |
|---|---|---|
| MCP (gp25) | 2-fold | B, C |
| | 3-fold | D |
| | 5-fold | G |
| | pseudo-6-fold | A-F |
| CP (gp26) | 2-fold | N |
| | 3-fold | J, K |
| | 5-fold | L, M |
| | pseudo 6-fold | 2x: H, I, J, K, N<br>1x: L, M |

**Table S4 |** Names and sequences for primers used in this study.

| Name | Sequence | Direction | Hybridisation site |
|---|---|---|---|
| ORF18_F | acgttctagacATGAAACTGGTTTCGATGAAGAAG | Forward | 5’-terminus of ΦXacm4-11 ORF18 |
| ORF18_R | acgtcccgggcATCCTTCATGCTCGGGTC | Reverse | 3’-terminus of ΦXacm4-11 ORF18 |
| ORF19_F | acgttctagacATGAGCAAATACGAATACACCC | Forward | 5’-terminus of ΦXacm4-11 ORF19 |
| ORF19_R | acgtcccgggcGTCCTGGCTGTTGGCG | Reverse | 3’-terminus of ΦXacm4-11 ORF19 |
| ORF20_F | acgttctagacATGTGCAGCAGCGCC | Forward | 5’-terminus of ΦXacm4-11 ORF20 |
| ORF20_R | acgtcccgggcCATCCCCAGCGCGG | Reverse | 3’-terminus of ΦXacm4-11 ORF20 |
| ORF22_F | acgttctagacATGGAAATCGAAGCC | Forward | 5’-terminus of ΦXacm4-11 ORF22 |
| ORF22_R | acgtcccgggcTGCGGCTTGCC | Reverse | 3’-terminus of ΦXacm4-11 ORF22 |
| ORF25_F | acgttctagacATGGCAATCCTCGCCAAC | Forward | 5’-terminus of ΦXacm4-11 ORF25 |
| ORF25_R | acgtcccgggcGACGACGACGCGGG | Reverse | 3’-terminus of ΦXacm4-11 ORF25 |
| ORF26_F | acgttctagacATGATCCTCGATCAGCTGC | Forward | 5’-terminus of ΦXacm4-11 ORF26 |
| ORF26_R | acgtcccgggcGTTGCGAGCCTGCGG | Reverse | 3’-terminus of ΦXacm4-11 ORF26 |
| ORF31_R | acgtcccgggcTCCCTTGCCCCCAGC | Reverse | 3’-terminus of ΦXacm4-11 ORF31 |
| ORF36_F | acgttctagacATGACCGTCTCCGCCAAC | Forward | 5’-terminus of ΦXacm4-11 ORF36 |
| ORF39_R | acgtcccgggcTGCGTTCGGACGCAG | Reverse | 3’-terminus of ΦXacm4-11 ORF39 |
