## Supplementary Movie1 Legend for "Cryo-EM structure analysis of phage ΦXacm4-11 that infects the phytopathogen *Xanthomonas citri*"

**Movie S1 | Clipping view of the portal-adaptor insertion at the special vertex.** Stepwise clipping of the portal-adaptor region at the special vertex of the capsid, illustrating how the tail complex is inserted into the major capsid protein shell. Lateral and top views are shown to provide complementary perspectives of the interface. The clipping progresses between two defined positions (indicated within the movie), allowing visualization of the structural transition. This movie complements **Figure S4**.
